# Extracellular Matrix Modulates Outgrowth Dynamics in Ovarian Cancer

**DOI:** 10.1101/2022.01.30.478322

**Authors:** Sarah Alshehri, Tonja Pavlovič, Sadaf Farsinejad, Panteha Behboodi, Li Quan, Daniel Centeno, Douglas Kung, Marta Rezler, Woo Lee, Piotr Jasiński, Elżbieta Dziabaszewska, Ewa Nowak-Markwitz, Dilhan Kalyon, Mikołaj P. Zaborowski, Marcin Iwanicki

## Abstract

Ovarian carcinoma (OC) forms outgrowths that extend from the outer surface of an afflicted organ into the peritoneum. OC outgrowth formation is poorly understood because there is limited availability of OC cell culture models to examine the behavior of cell assemblies that form outgrowths. Prompted by immunochemical evaluation of extracellular matrix (ECM) components, laminin γ1 and collagens, in human tissues representing untreated and chemotherapy-recovered OC, we developed laminin- and collagen-rich ECM-reconstituted cell culture models amenable to studies of cell clusters that can form outgrowths. We demonstrate that ECM promotes outgrowth formation in fallopian tube non-ciliated epithelial cells (FNE) expressing mutant p53-R175H and various OC cell lines. Outgrowths were initiated by cells that had undergone outward translocation and, upon mechanical detachment, could intercalate into mesothelial cell monolayers. Electron microscopy, optical coherence tomography (OCT), and small amplitude oscillatory shear experiments revealed that high ECM concentration increased ECM fibrous network thickness and led to high shear elasticity in the ECM environment. These physical characteristics were associated with the suppression of outgrowths. A culture environment with low ECM concentration mimicked viscoelasticity of malignant peritoneal fluids (ascites) and supported cell proliferation, cell translocation, and outgrowth formation. These results highlight the importance of ECM microenvironments in modulating OC growth and could provide an additional explanation of why primary and recurrent ovarian tumors form outgrowths that protrude into the peritoneal cavity.

## INTRODUCTION

Ovarian carcinoma (OC) can arise from the outer mucosa of the epithelial cell surfaces covering the reproductive organs including the endometrium,^[1]^ ovary,^[2]^ and fallopian tubes.^[3]^ Within malignant tissue, transformed epithelial cells can form outgrowths capable of protruding away from the basement membrane and sub-epithelial stroma into the peritoneal cavity.^[4]^ Outgrowths can detach, disseminate within malignant peritoneal fluid (ascites), superficially attach, and intercalate into the mesothelium, contributing to the disease progression.^[5, 6]^ The ascites contain ECM molecules^[7]^ and ECM is also enriched within cell-cell junctions of detached tumor outgrowths.^[8]^ Thus, investigating the contribution of ECM to processes associated with OC outgrowth formation would provide invaluable information about the mechanisms of disease progression.

ECM is a molecular scaffold that is rich in numerous glycosylated proteins including laminin γ1^[9]^ and collagens^[10]^ which, through interaction with cell surface receptors,^[11]^ provide mechanical^[12–15]^ and biochemical^[16, 17]^ cues competent in regulating tissue development^[18, 19]^ and malignant progression.^[20, 21]^ Recent histologic examination of OC outgrowths protruding from the fallopian tube demonstrates enrichment of laminin γ1 within tumor cells capable of forming outgrowths.^[22]^ Consistent with these findings, it was previously reported^[8]^ that detached human fallopian tube non-ciliated epithelial (FNE) cells expressing various p53 mutations deposited ECM molecules supportive of both tumor survival and cell-ECM-cell adhesion. A recent study using second harmonic visualization of outgrowths arising in the fallopian tube revealed the presence of additional ECM components, collagens.^[23]^ Collagen molecules were found to form fibrillar and wavy networks beneath basal surfaces of transformed epithelium. Taken together, these data provoked questions of whether constituents of the tumor microenvironment, such as ECM, regulate the dynamics of outgrowth-forming cell clusters.

Reconstitution of cell cultures with Matrigel® (MG), a basement membrane extract rich in both laminin and collagen, has become a standard approach^[24]^ in studying the role of ECM in cellular growth,^[25]^ migration,^[26]^ and death.^[27]^ Recent studies^[28–31]^ employed MG to grow OC cells to evaluate their response to treatments in the context of three-dimensional (3D) spheroid cultures, which resemble malignant tissue organization more closely than two-dimensional (2D) cell cultures. While these studies provided a static view of tumor cells with ECM reconstitution, the contribution of ECM to the dynamic process of OC growth remains unknown.

In this report, we have performed immunochemical evaluations of laminin γ1 and collagens in OC representative of disease progression before and after chemotherapy. Similar to OC tumorigenesis from the fallopian tube,^[22]^ we found that laminin γ1 was associated with tumor cells, whereas collagens formed fibrillar networks within the surrounding microenvironment. These studies raised a question of whether ECM deposition within tumor cells and adjacent microenvironment contribute to OC outgrowth dynamics. To address this question, we deployed laminin- and collagen-rich (MG) ECM-reconstituted suspended or adhered cell culture models. To study the dynamics of OC cell clusters, live-cell imaging and quantitative image processing of cell proliferation and translocation were employed. We found that ECM reconstitution of suspended cell clusters induced outgrowths in FNE cells expressing mutant p53-R175H, the cells-of-origin for a significant subset of OCs, as well as other genetically distinct and well-established OC cell lines. Live-cell imaging revealed that outgrowths were initiated by cell clusters that had undergone outward translocation and, upon mechanical detachment, could intercalate into mesothelial cell monolayers – the most frequent physiologic sites of metastasis for OC. Increased ECM concentration altered the thickness of the fibrous network and the viscoelastic characteristics of the microenvironment, which led to the suppression of cell proliferation, directional cell translocation, and formation of outgrowths. Furthermore, ECM-reconstituted cultures that supported outgrowths, cell proliferation, and cell translocation mimicked the viscoelastic properties of peritoneal fluids (ascites).

Our results are consistent with a model whereby low-elasticity ECM microenvironments support cell proliferation and translocation of cell clusters that form outgrowths. Our data provide additional explanation of why primary and recurrent OCs favor forming outgrowths that protrude into peritoneal cavity, as opposed to breaching underlying collagen-dense tissues.

## RESULTS

### OC tumor outgrowths are associated with distinct ECM deposition

Detached OC cell clusters deposit ECM on cell surfaces to support survival and cell-ECM- and cell-cell adhesion.^[8]^ Pathologic^[4]^ and genomic^[32]^ examination of tumors indicate that the sources of detached OC cell assemblies are malignant outgrowths that can detach and disseminate into the peritoneal space from the surface of the fallopian tubes, ovaries, and the omentum. To examine whether OC outgrowths are associated with ECM deposition, we used combination of immunohistochemistry (IHC) for laminin γ1 and picrosirius red staining for collagen.^[34]^ This approach revealed that cell clusters, which were positive for OC marker PAX8, grew outward from the human ovary surface (**Figure 1A**), or protruded from the surface of the omentum in patient-derived xenograft (PDX) models of OC (**Figure 1B**). These OC outgrowths contained laminin γ1 (**Figure 1C, D**) and, to a much lesser degree, collagens (**Figure 1E, F**). In the same tissue sections, we observed that the adjacent extra-tumoral connective space exhibited low laminin γ1 levels but contained dense fibrillar collagen networks **(Figure 1C-F**). Quantitative image analysis of laminin γ1 and collagen expression revealed differential patterns of localization. Laminin γ1 was enriched within tumors (**Figure 1G, I**), whereas collagens were enriched within the extra-tumoral space (**Figure 1H, J**). Taken together, these results demonstrate that OC outgrowths are associated with at least two ECM microenvironment factors, laminin γ1 and collagens.

**Figure 1.**
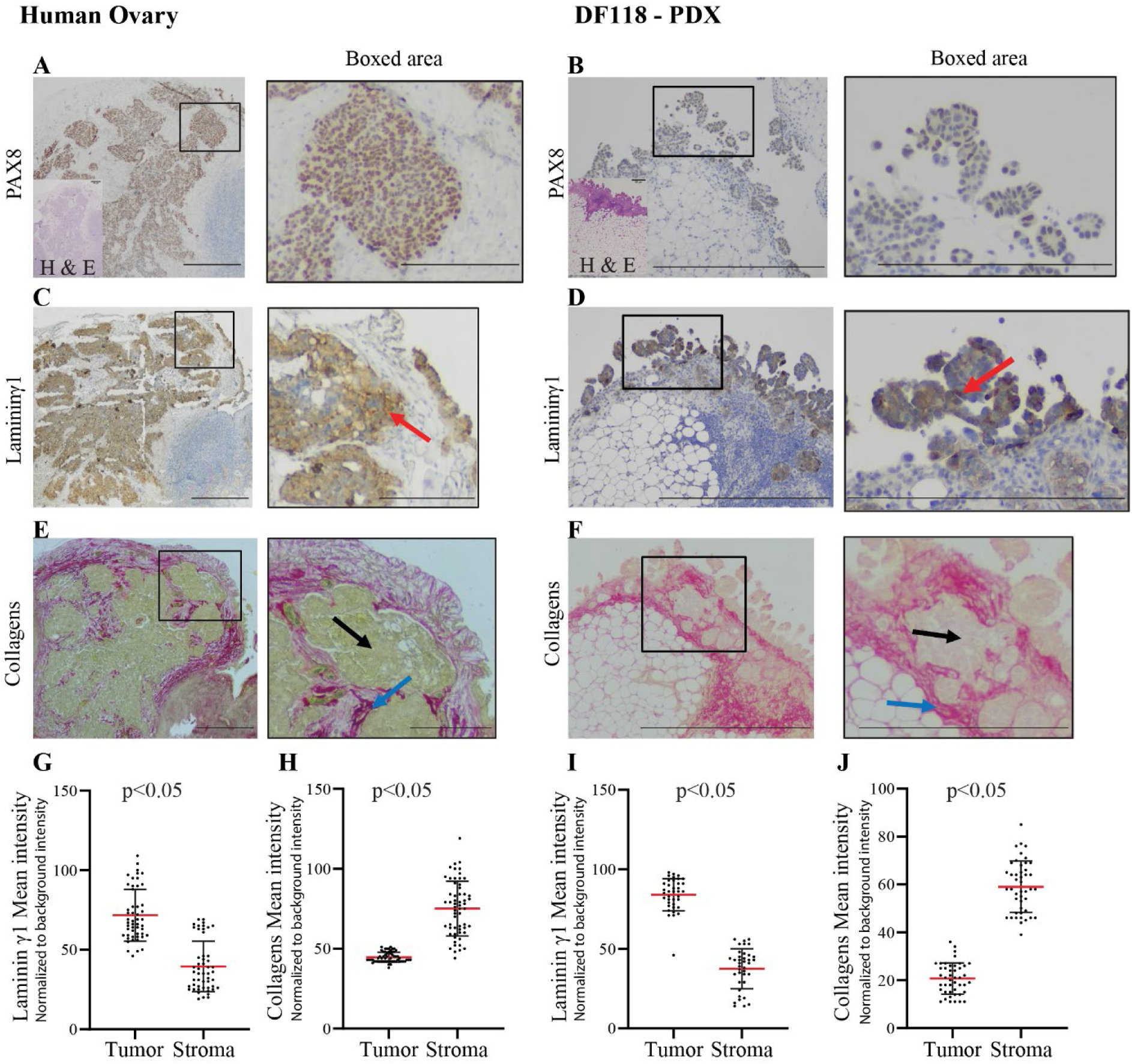
Immuno and chemical evaluation of laminin γ1 and collagen. (**A, B**) PAX8-positive OC outgrowths, and PAX8-negative surrounding connective tissue. Representative images of laminin γ1 (**C, D**), and collagen (**E, F**) expression in tumors protruding from human ovary (**C, E**), or in xenograft experiments, from the surface of the omentum (**D, F**). Red arrows point to the deposition of laminin γ1. Blue arrows indicate fibrillar collagen structures within the tumor’s connective tissue, whereas black arrows represent collagen deposition within tumors. Quantification of laminin γ1 (**G, I**) and collagen (**H, J**) expression in PAX8-positive tumor outgrowths and PAX8- negative surrounding tissue. Each data point represents one region of interest (ROI) within tumor outgrowth or surrounding stroma. In each group, G: 51, H: 59, I: 38, and J: 45 ROIs/ condition (tumor and stroma). All data points are shown with bars indicating mean ± SD. An unpaired, two-tailed, non-parametric Mann-Whitney test was used to examine statistical differences between data sets. Intensity values are normalized to background level (area without tissue) after RGB image conversion to 8-bit gray scale (0-255 pixels range).^[33]^ Bars are 100 µm and 50 µm in boxed area.

### Laminin γ1 and collagens are associated with OC tumors before and after chemotherapy

OC patients often develop recurrent disease, which is characterized by the continuous evolution of outgrowths and the presence of detached tumor clusters in the peritoneal cavity.^[7]^ These clinical observations prompted us to examine whether chemotherapy-recovered tumors are associated with laminin γ1 and a collagen-rich ECM microenvironment. ECM deposition was evaluated in PAX8-positive primary tumors in the ovary (**Figure 2A**) and matching omental metastases (**Figure 2B-C**), representing tumors before and after neoadjuvant chemotherapy. Examination of these tissue samples revealed laminin γ1 and collagen deposition before and after chemotherapy (**Figure 2D-E**), suggesting that OC after chemotherapy is associated with the presence of laminin γ1 and collagens. These findings support the idea that laminin and collagen-rich ECM could contribute to outgrowth dynamics of primary and recurrent tumors.

**Figure 2.**
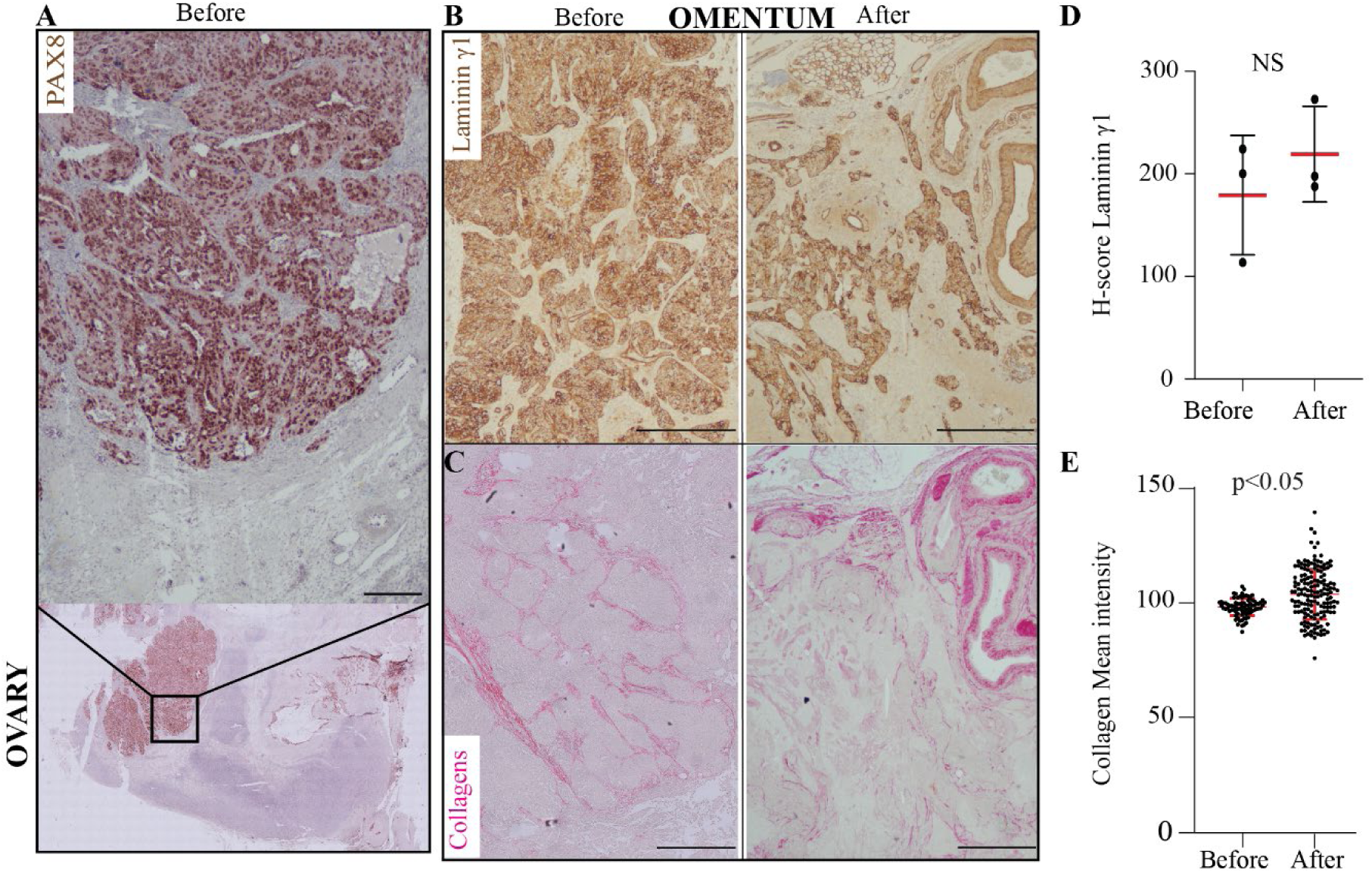
OC before and after chemotherapy is associated with spatially distinct laminin γ1 and collagen ECM microenvironments. (**A**) Ηematoxylin and PAX8 stain of high-grade serous ovarian cancer (HGSOC) growing within the ovary before chemotherapy; bar is 100μm. (**B**) Laminin γ1 and (**C**) collagen deposition in matched omental tissue of HGSOC before and after chemotherapy; bar is 500μm. (**D**) H-score quantification of laminin γ1 positivity in matched tissue samples representing HGSOC before and after taxane-platin therapy. Tissues were analyzed from three patients (n=3). Wilcoxon sum rank test was used to compare the difference between the groups, NS= Not-Statistically different. Data points are presented as mean ± SD. (**E**) Quantification of collagen expression in omental metastases before and after chemotherapy. Each data point represents one region of interest (ROI); Before: 82 & After: 160 ROIs. All data points are shown with bars indicating mean ± SD. An unpaired, two-tailed, parametric student t-test with Welch’s correction was used to examine statistical differences between data sets. Intensity values are normalized to background level (area without tissue) after RGB image conversion to 8-bit grayscale (0-255 pixels range).^[33]^

### ECM reconstitution in OC spheroids promotes outgrowths

Laminins (including laminin γ1) and collagens (including collagen IV) represent major scaffolding components of the basement membrane^[35–37]^ and of the reconstituted basement membrane extract, Matrigel® (MG).^[37]^ Thus, to mimic a laminin γ1 and collagen microenvironment associated with OC, we tested whether addition of MG would support formation of OC outgrowths in a suspended spheroid cell culture model. We initially examined effects of MG addition on outgrowth formation in normal fallopian tube non-ciliated epithelial cells expressing mutant p53 R175H (FNE-m-p53). We selected this cell line, because (1) non-ciliated fallopian tube cells, with mutations in the *TP53* gene, are thought to be precursor cells of OC,^[3]^ and (2) we have recently demonstrated, using this cell line, that expression of m-p53 in FNE cells induces acquisition of transformed phenotypes associated with OC progression, such as survival of suspended spheroids and mesothelial intercalation.^[8]^ Here, we examined the outgrowth-forming capabilities of FNE-m-p53 or FNE cells transduced with an empty control vector in an OC spheroid model. To obtain OC spheroids, we clustered OC cells in low-adhesion wells and allowed them to grow in suspension for 24 hours (**Figure 3A**). To mimic the laminin and collagen rich ECM microenvironment, we then embedded the OC spheroids in media containing 2% MG (**Figure 3A**). This ECM reconstitution condition of 2% MG was selected based on previous reports that 2% MG supported the growth of breast and ovarian spheroid cultures plated on MG layers covering flat culture surfaces.^[27]^ Live-cell microscopy revealed that FNE-m-p53, but not wild-type FNE, significantly expanded and formed outgrowths – cell assemblies that, over time, began to extend beyond the main spheroid structure (**Figure 3B-C, and MOVIE 1**). We observed that FNE-m-p53 spheroids in 2% MG formed outgrowths with an average frequency of 30%, whereas no outgrowths were detected in spheroids composed of control FNE cells in 2% MG, or FNE-m-p53 spheroids lacking MG embedding (**Figure 3D**). These data are consistent with the implicated role of ECM deposition in supporting outgrowth development (**Figure 1 and Figure 2**).

**Figure 3.**
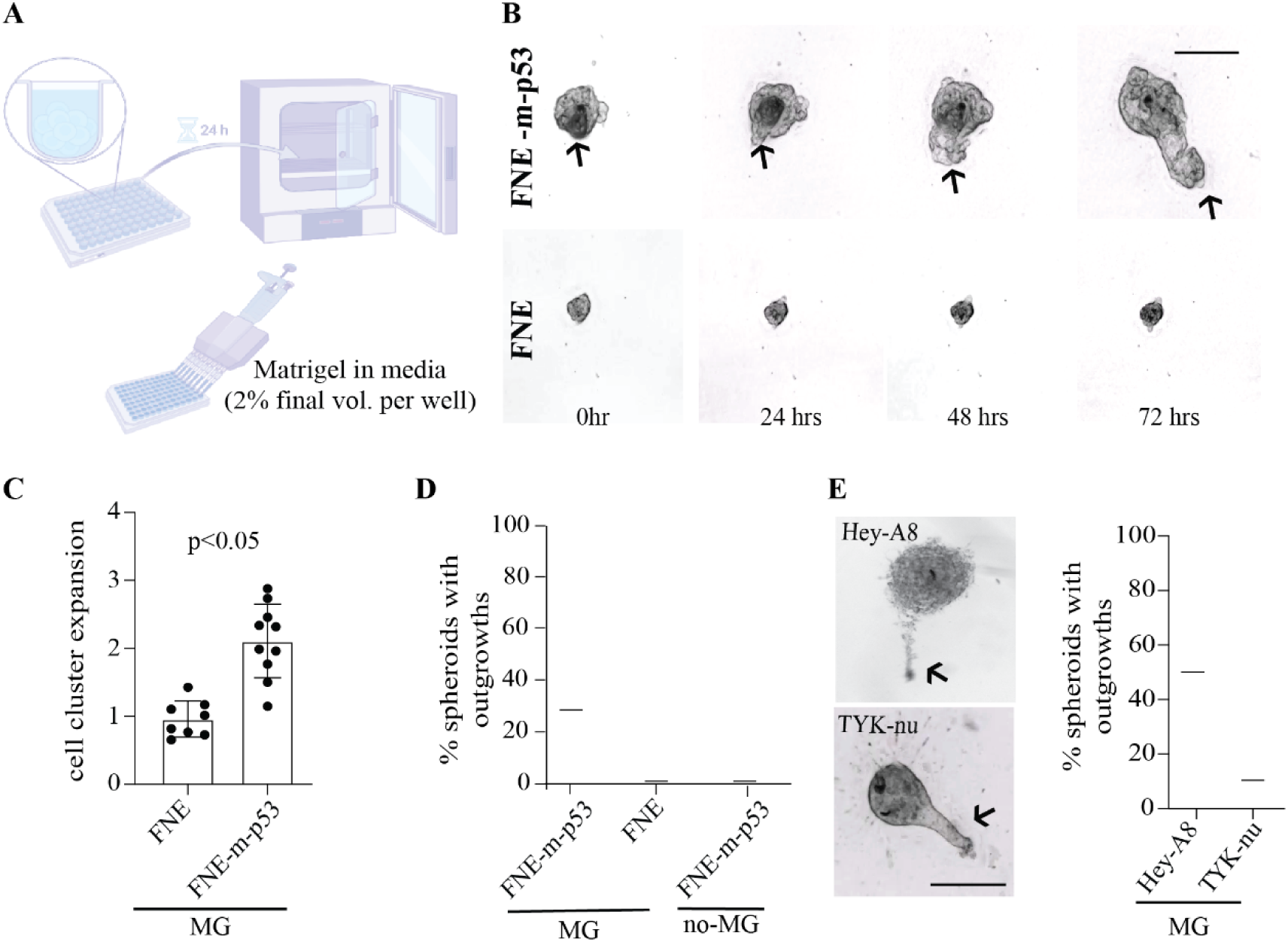
Laminin- and collagen-rich ECM reconstitution stimulates outgrowths in suspended cultures of various OC cells. (**A**) Graphical representation of the assay design to study outgrowth dynamics. (**B**) Representative bright-field images of outgrowth formation in FNE cells expressing plasmid containing m-p53R175H (FNE-m- p53), or control plasmid, and cultured as suspended clusters reconstituted with 2% MG. Cell structures were cultured for 7 days before imaging and subsequently followed for an additional 72 hrs. (**C**) Quantification of 3D structure (spheroid) expansion, where each dot represents fold change in area, over 72 hrs of filming time, in a spheroid. Total spheroids: 8 for FNE & 10 for FNE-m-p53. All data points are shown with bars indicating mean ± SD. An unpaired, two-tailed, parametric student t-test with Welch’s correction was used to examine statistical differences between data sets. (**D**) Quantification of cell clusters with outgrowths in FNE-m-p53 or FNE cells. Bar represents an average percentage of outgrowth formation with at least 30 cell clusters analyzed. (**E**) Representative bright-field images of outgrowths in Hey-A8 and TYK-nu ovarian cancer cell lines, and quantification of outgrowths in Hey-A8 or TYK-nu. Bars represent an average percentage of outgrowth formation in three independent experiments with 20-30 spheroids scored per experiment. Scale bars are 200 μm.

Intraperitoneal dissemination is a common feature among different subtypes of OC, including serous and non-serous subtypes.^[6]^ We therefore wanted to evaluate whether multiple OC cell lines, representing a diverse variety of OC subtypes, can form similar outgrowths when cultured as suspended spheroids embedded in 2% MG. We examined OC cell lines Hey-A8 and TYK-nu, representing non-serous and likely serous OC models, as defined by Domcke et al.^[38]^ Similar to FNE-m-p53 spheroids, reconstitution of Hey-A8 (**MOVIE 2**) or TYK-nu (**MOVIE 3**) spheroids with 2% MG prompted outgrowth formation (**Figure 3E**). The average percentage of outgrowth formation among Hey-A8 and TYK-nu spheroids varied from 50% to 10%, respectively (**Figure 3E**). Outgrowth formation in spheroids observed in different OC cell lines, but not normal FNE cells, suggests that ECM collaborates with transformation to support outgrowths.

### Detached outgrowths clear mesothelial monolayers

OC outgrowths can ultimately detach from the transformed tissue, transit into the fluids of the abdominal cavity, and intercalate into mesothelial surfaces of other abdominal organs, such as the omentum.^[1]^ To assess whether OC outgrowths, evoked by ECM reconstitution, possess the ability to intercalate into mesothelial cell layers, we utilized Hey-A8 spheroids embedded in 2% MG due to their tendency to form long protruding outgrowths that are amenable to dissociation under a dissecting microscope. We mechanically separated Hey-A8 outgrowths from the spheroid structure and co-cultured both entities on top of a mesothelial cell monolayer expressing green fluorescent protein (GFP) (**Figure 4A**). To examine the ability of spheroids and their corresponding outgrowths to invade mesothelial cell layers, we used a well-established mesothelial clearance assay,^[40]^ which relies on measuring the area where initially adherent mesothelial cells are displaced by the introduction of OC spheroids.^[39]^

**Figure 4.**
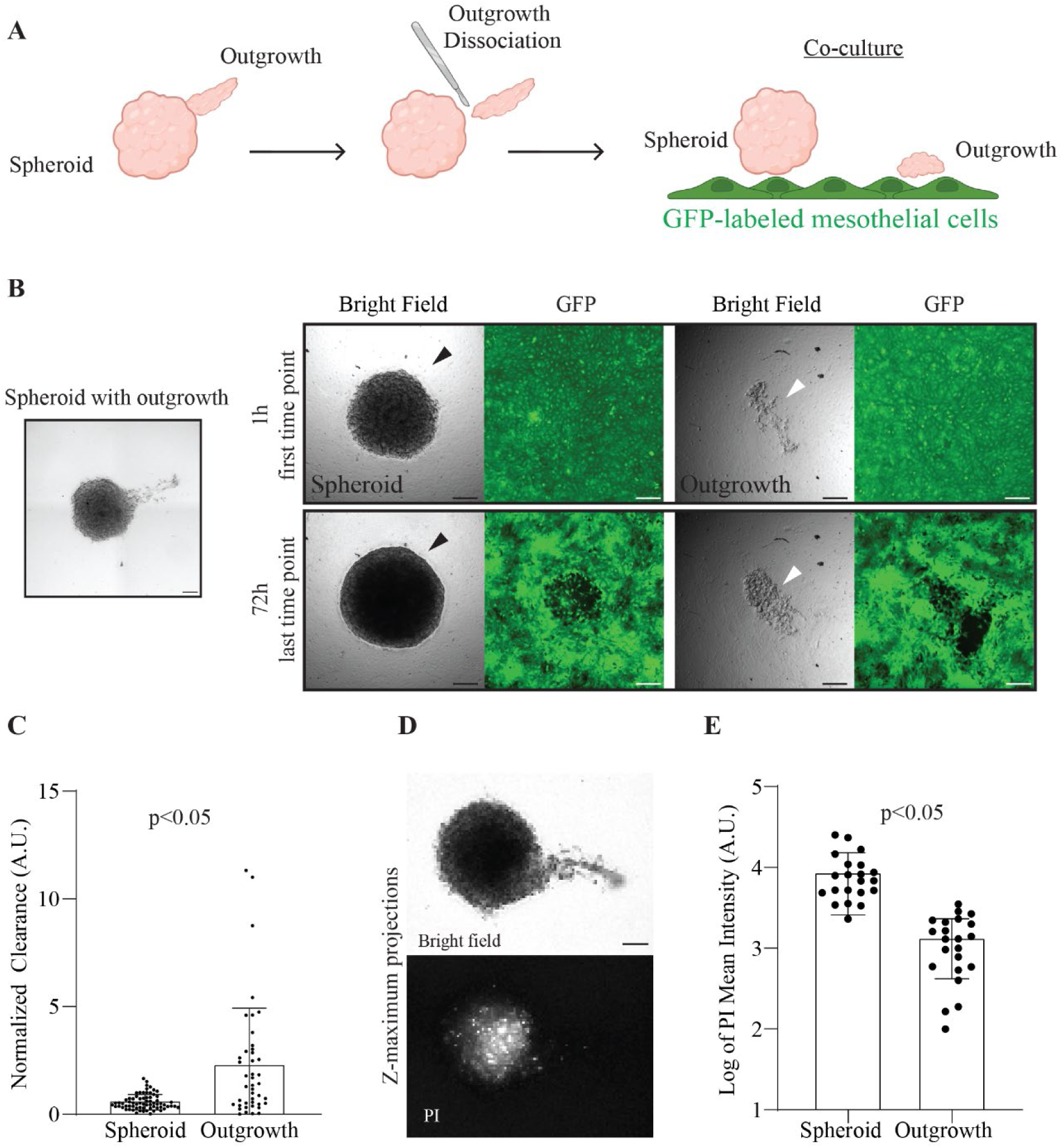
Detached outgrowths clear mesothelial monolayers. (**A**) Schematic representation of Hey-A8 outgrowth detachment and co-culture with mesothelial cells expressing GFP. Created with BioRender.com. (**B**) Representative bright-field image of a spheroid before outgrowth detachment, followed by representative bright- field and fluorescent images of mesothelial clearance by Hey-A8 spheroid (black arrow) or detached outgrowth (white arrows); scale bar is 200 μm. (**C**) Quantification of mesothelial clearance by Hey-A8 spheroids and detached outgrowths.^[39]^ Each data point represents a spheroid or outgrowth from two independent experiments from 69 spheroids and 43 outgrowths. (**D**) Representative bright-field and corresponding fluorescent maximum projection images of a Hey-A8 spheroid forming an outgrowth and treated with propidium iodide (PI) to spatially visualize cell death within the structure; scale bar is 200 μm. (**E**) Quantification of PI incorporation by spheroids or outgrowths. Twenty-one Hey-A8 cell structures were analyzed. In C and E all data points are shown with bars indicating mean ± SD, and an unpaired, two-tailed, non-parametric, Mann-Whitney test was used to examine statistical differences between data sets.

The formation of the cleared area is visualized by the time-dependent exclusion of GFP fluorescence, which is representative of mesothelial cell displacement from beneath the intercalating OC spheroid. Using this displacement assay, we were able to observe that detached outgrowths induced more mesothelial cell clearance compared to their corresponding OC spheroid clusters (**Figure 4B-C**). This observation could be due to spatial variation in viability between OC cells located within the outgrowth and cells within the spheroid. Spatial examination of cell viability using propidium iodide (PI), a fluorescent dye permeable only to dead cells, revealed a greater quantity of dead cells localized to the core of the OC spheroids relative to the outgrowths, which demonstrated minimal PI incorporation and thus increased cell viability (**Figure 4D-E**). These observations are consistent with the hypothesis that OC cell assemblies capable of forming outgrowths are viable and can efficiently intercalate into mesothelial cell layers upon detachment.

### Increased ECM inhibits formation of outgrowths

As we have so far observed that ECM reconstitution of OC spheroids was able to induce formation of cell outgrowths that were able to intercalate into mesothelial cell layers, we wanted to further investigate the role of ECM in the regulation of OC cell outgrowths. Motivated by our initial observation indicating that OC prefers to grow away from collagen dense areas (**Figure 1**), we examined whether manipulation of ECM concentrations affect outgrowth formation. We compared the outgrowth formation capability of FNE-m-p53 and Hey-A8 spheroids reconstituted with 2% or 25% MG. As expected, reconstitution of both cell types with 2% MG induced outgrowth formations. In contrast, 25% MG culturing conditions significantly attenuated the development of outgrowths (**Figure 5A-B**), indicating that increasing ECM concentrations can suppress outgrowths. In addition, we observed that under 25% MG conditions, both FNE-m-p53 and Hey-A8 spheroids appeared to be significantly smaller when compared to 2% MG reconstitution conditions (**S. Figure 1A-B**). In comparison to FNE-m-p53 spheroids, Hey-A8 spheroids cultured in 25% MG also appeared to be less compacted, coarse, and more disorganized (**Figure 5B, inset**). This may suggest that elevating ECM concentrations can have divergent effects on different OC cell lines. In the case of Hey- A8 spheroids, increased concentrations of ECM may have disrupted cell-cell adhesions, as suggested by the decrease in N-cadherin staining at the edge of the spheroids (**S. Figure 2**). Decrease in spheroid size due to an increase in ECM concentrations indicated the possibility that increasing ECM deposition could affect cell proliferation.

**Figure 5.**
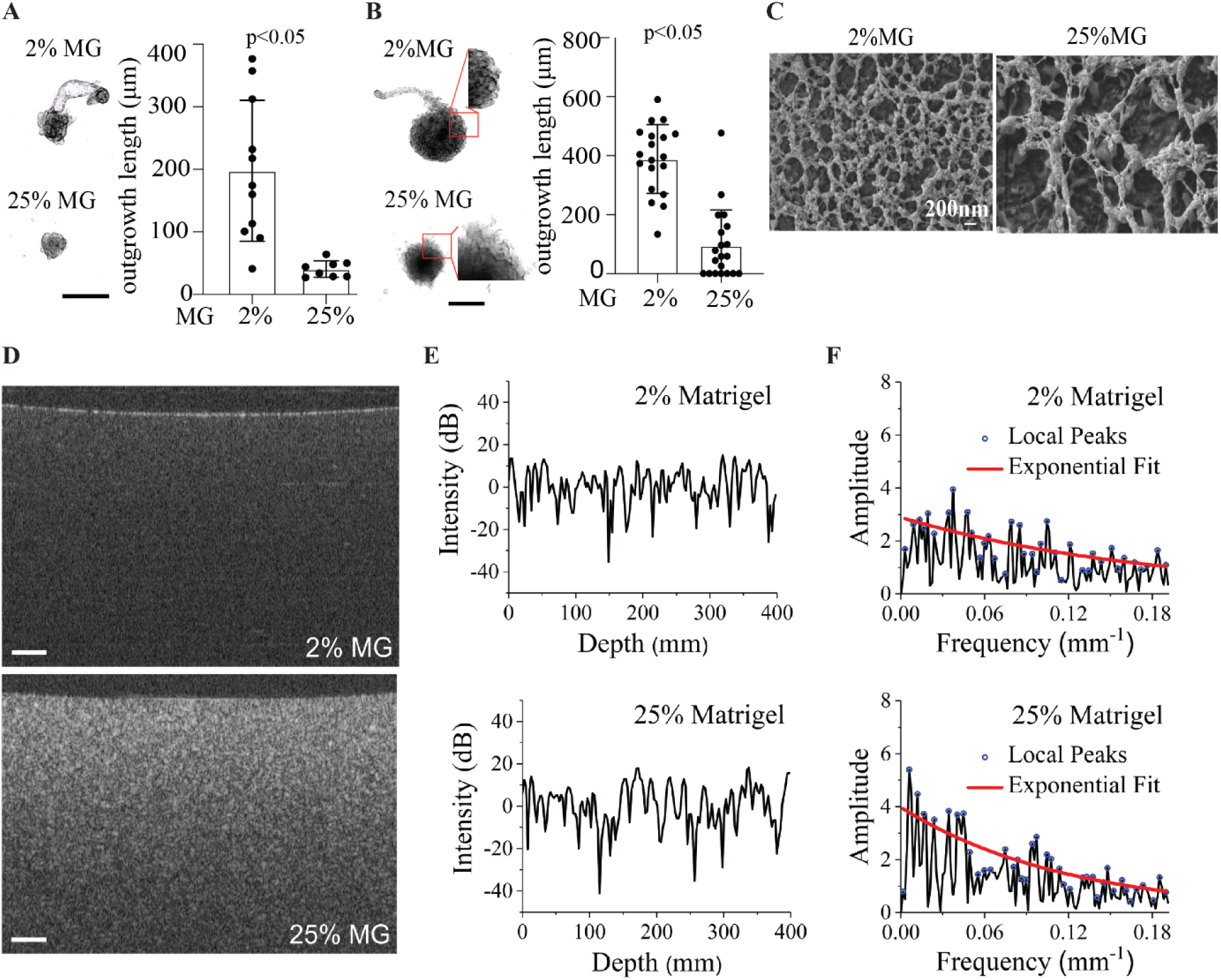
Elevation of ECM concentration inhibits outgrowths. Representative bright-field maximum Z-projection images of (**A**) FNE-m-p53 spheroids reconstituted with 2% or 25% MG; scale bar is 250 μm. (**B**) Hey- A8 spheroids reconstituted with 2% or 25% MG; scale bar is 500 μm. Spheroids were grown for 10 days, and the images show the final time point. Dot plots represent the distribution of outgrowth lengths from individual spheroids reconstituted with 2% or 25% MG. All data are shown with bars indicating mean ± SD. An unpaired, two-tailed, parametric student t-test with Welch’s correction was used to examine statistical differences between data sets. (**C**) Representative electron scanning micrographs representing 2% and 25% MG. (**D**) Representative OCT B-scan images of 2% and 25% MG; scale bars are 100 µm. (**E**) Representative OCT depth-resolved intensity profiles where the slopes from optical attenuation are removed. (**F**) Amplitude spectra of the spatial frequency corresponding to the panel -E- show relatively stronger low-frequency components from 25% MG, and exponential fit of local peaks indicating a faster decay rate in 25% MG.

To explore this possibility, we took advantage of a 2D cell culture assay in which the cell monolayer is overlaid with MG,^[41]^ thereby partially mimicking conditions of ECM encapsulation (**S. Figure 3A**). We integrated this assay with live-cell imaging of FNE-m-p53 and Hey-A8 cells stably expressing GFP or monomeric Kusabira-Orange2 (mKO2), respectively. FNE-m-p53-GFP or Hey-A8-mKO2 cells cultured with 25% MG significantly decreased cell proliferation (**S. Figure 3B-C, MOVIE 4 and 5**). These results were additionally supported by data indicating that 25% MG reconstitution of Hey-A8 spheroids significantly reduced the number of cells undergoing active DNA replication, as assessed by the 5-ethynyl-2’-deoxyuridine (EdU) incorporation assay (**S. Figure 3D**). We selected Hey-A8 spheroids to examine EdU incorporation due to the formation of significantly larger spheroid structures when compared to FNE-m-p53 cells (**S. Figure 1A-B**), enabling robust EdU incorporation analysis. Taken together, these results suggest that ECM play a significant role in the OC cell environment and can modulate OC cell proliferation and outgrowth formation.

### Increased ECM leads to the formation of thick fibrillar networks

The phenotypic influence of ECM concentrations on outgrowth formation and cell proliferation (**Figure 5A-B, and S. Figure 3**), combined with the previous IHC examination of tumors, which indicate that OC tumors can form protruding outgrowths away from thick collagen fibrous network associated with tumor-surrounding tissue (**Figure 1**), motivated us to examine whether higher ECM concentrations exhibit differences in fibrous network thickness. Scanning electron microscopy (SEM) of MG revealed qualitative differences between ECM networks formed in 2% and 25% MG (**Figure 5C**); network fibers formed in 25% MG appeared to be thicker and more spaced out (larger spaces between fibers) as compared to network fibers formed in 2% MG. For quantitative evaluation of ECM fibers, we imaged the MG samples using optical coherence tomography (OCT), where the endogenous optical contrast describes the microstructure of the sample. The OCT B-scan images revealed a higher contrast from the ECM contents with larger high-intensity clusters in the 25% MG condition compared with the 2% MG condition (**Figure 5D**). Also, the representative OCT depth-resolved intensity profiles where the slopes from optical attenuation are removed, show a larger spatial period of intensity fluctuations in the 25% MG (**Figure 5E**), which can be clearly seen with relatively more low-frequency components from the corresponding amplitude spectra (**Figure 5F**). This feature in the spatial frequency was quantified with the decay coefficient from an exponential fit of the local peaks in the amplitude spectra (**Figure 5F**), where a higher exponential decay coefficient in 25% MG indicates relatively stronger low-frequency components, suggesting thicker ECM fibers and/or larger space between ECM fibers in 25% MG.

### ECM modulates viscoelasticity of OC culture microenvironments

Based on data demonstrating formation of thicker, spatially distributed – ECM fiber networks in 25% MG cultures, and previous reports indicating positive correlation between MG concentration and elasticity,^[42]^ we wondered whether increasing the concentration of ECM in our model would affect the cell culture’s viscoelasticity e.g., changes to time dependent elasticity and the viscous behavior of MG in the cell culture.^[43]^ We characterized the linear viscoelastic material functions of cell culture media containing 2% or 25% MG. This analysis revealed that in culture media supplemented with 25% MG, the loss modulus (G”) values representing viscous behavior, which are indicative of the energy dissipated as heat when the culture media is deformed under small-amplitude oscillatory shearing, were within a narrow range of 0.5 to 0.9 Pa in the frequency range of 0.1 to 10 s⁻¹. The loss modulus values were thus not sensitive to the frequency of rotation (**Figure 6A**). Under similar conditions, the storage modulus (G’) values representing elasticity i.e., indicative of the energy stored as elastic energy during the deformation of the culture media, were within the range of 3.5 to 4.5 Pa. The storage modulus (G’) values were also insensitive to the frequency of rotation (**Figure 6A**).

**Figure 6.**
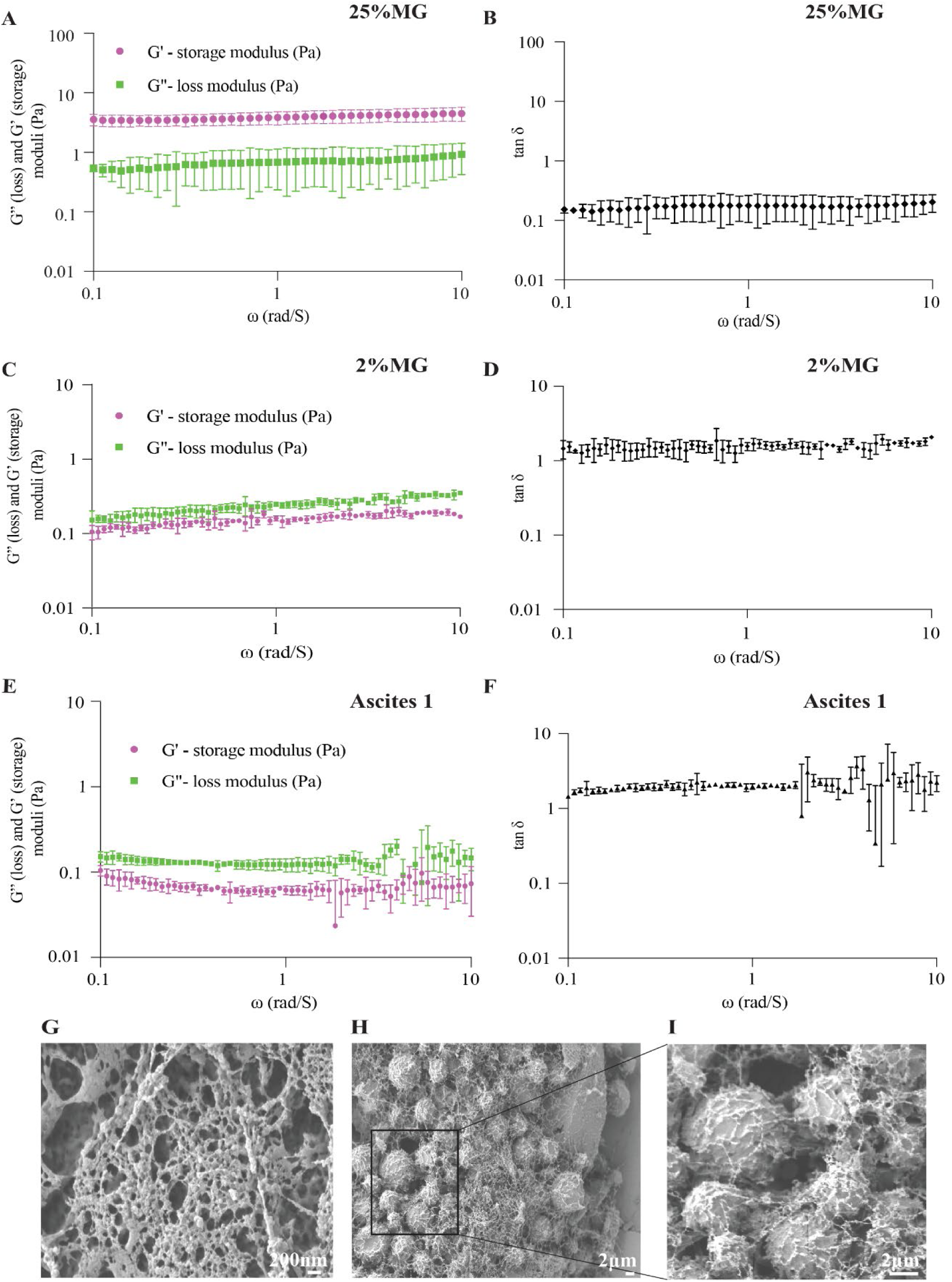
ECM modulates viscoelastic properties in the OC culture microenvironment. Measurements of storage (elastic) and loss (viscous) moduli in OC cell culture media reconstituted with 2% or 25% of MG (**A, C**); or in ascites isolated from OC patients with a relapsed disease (**E**). (**B, D, F**) Graphs represent calculated tan δ from values reported in A, C & E. (**G, H, I**) Scanning electron micrographs represent the ascitic fibrous network surrounding tumor cells. Each data point represents the mean and SD from triplicate measurements with a total of 41 (25% MG), 91 (2% MG) & 91 (ascites) measurements.

The loss modulus (G”) values of samples cultured in 2% MG ranged between 0.05 to 0.1 Pa while the storage modulus (G’) values were between 0.09 to 0.3 Pa (Figure 6C). The storage moduli of 25% MG reconstituted culture media were significantly higher than the loss moduli (G’ > G”), and the ratio of G’’/G’ (tan δ) was below 1 (Figure 6B). On the other hand, the 2% MG reconstitution condition produced tan δ values greater than 1 (Figure 6D), indicating a less elastic nature of the 2% versus the 25% MG. As expected, these results suggest that the elasticity of media containing 25% MG is significantly higher than that containing 2% MG. Measurements of loss and storage moduli of ascites isolated from three different OC patients exhibited a closer similarity to the values of G”, G’ and tan δ observed in the 2% MG reconstitution condition (Figure 6E-F and S. Figure 4B-C), indicating that ascites may mechanically appear as gels with lower elasticity. Although the study of ascitic viscoelasticity was limited to 3 patient samples that showed considerable G’ and G’’ variation (S. Figure 4B- C), all the samples exhibited a relatively low storage modulus when compared to 25% MG. Behavior of ascites as weak gels with low torque values could also account for the variability, because data points associated with very low rotation frequencies might be close to the limits the instrument’s sensitivity. Nevertheless, the similarity of viscoelastic properties between 2% MG and patient-derived ascites motivated us to evaluate whether ascites possess the ability to form extracellular fibrous networks. Scanning electron microscopy of ascitic fluid indicated the presence of structures resembling extracellular fiber networks (Figure 6G). Additionally, fibrous structures were associated with cells that were isolated along with ascites. Moreover, we have detected, using IHC, laminin and collagen fiber-like structures that were associated with detached OC clusters isolated from human or mouse ascites (**S. Figure 5**). Taken together, these results are consistent with the hypothesis that ECM microenvironments with low shear viscosity and elasticity, support OC outgrowth formation.

### Increased ECM suppresses directional translocation

Based on observations that outgrowths are initiated by cell assemblies that translocate outside the main spheroid structure (**Figure 3, MOVIE 1, 2, and 3**), and that increasing ECM concentration suppresses the formation of outgrowths, decreases spheroid size, and impedes cell proliferation (**Figure 5A-B, and S. Figures 1 and 3**), we hypothesized that high ECM concentrations also suppress cell translocation.

To track the movement of cells embedded in ECM, we overlaid FNE-m-p53-GFP and Hey- A8-mKO2 cells grown on a 2D surface with media containing 2% or 25% MG (**S. Figure 3A**). We analyzed the cell translocation dynamics of FNE-m-p53-GFP and Hey-A8-mKO2 cells overlaid with differing MG concentrations using an ImageJ-integrated plugin, TrackMate and measured the confinement ratio output, which represents the efficiency of cell translocation away from its original starting point (**Figure 7A**; see Materials and Methods for details). We observed that overlaying FNE-m-p53-GFP or Hey-A8-mKO2 cells with 2% MG resulted in a confinement ratio that was significantly larger than the confinement ratio of cells overlaid with 25% MG, indicating more efficient cell translocation (**Figure 7B-E**). Cell tracks appeared more persistent and directional in cells overlaid with 2% MG medium as compared to 25% MG medium (**Figure 7B, D, and MOVIE 6 and 7**), suggesting that an increase in ECM deposition can suppress directional cell translocation and is competent in anchoring cells to their starting point, as supported by the increased number of focal adhesions under 25%MG conditions (**S. Figure 6** and **S. Figure 7**). Furthermore, we observed that OC cells superficially attached to the outer surface of murine small intestine did not invade beyond smooth muscle actin positive stroma (**S. Figure 8**), supporting that some OC tumors can grow superficially and do not penetrate beyond stroma and dense ECM.

**Figure 7.**
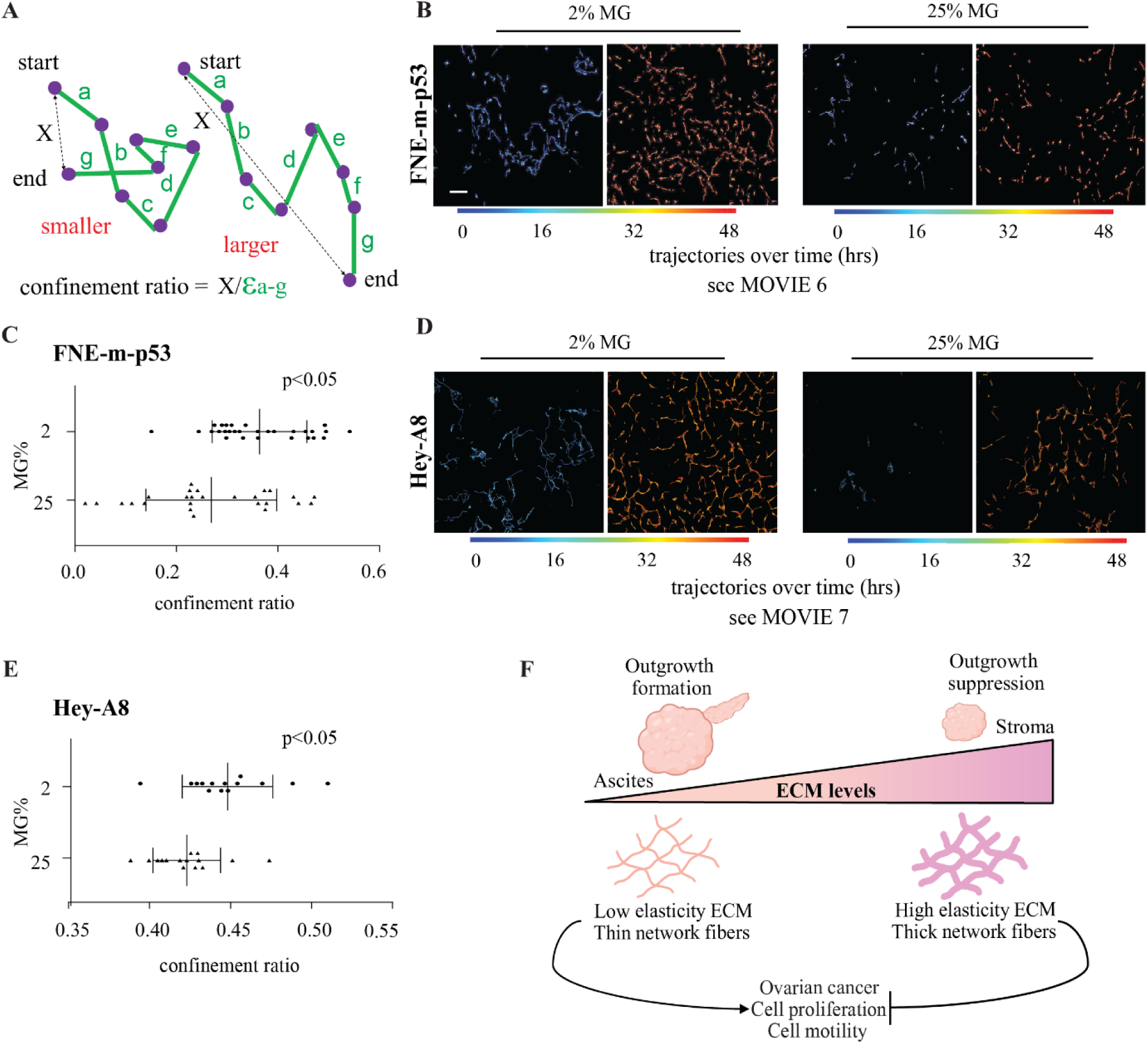
Elevation of ECM concentration suppresses directional cell translocation. (**A**) Graphical illustration of confinement ratio quantification. (**B**) Representative trajectory images of cell translocation within FNE-m-p53 or (**D**) Hey-A8 cell monolayers overlayed with 2% or 25% MG. Each line represents a single cell trajectory formed during 50 frames/time interval. Time zero corresponds to the first 50 frames; scale bar 100 μm. Distribution of confinement ratio within (**C**) FNE-m-p53 or (**E**) Hey-A8 cell monolayers overlayed with various concentrations of MG. Each dot represents one field of view that contained 20-100 cells/ condition (2% vs. 25% MG). All data points are shown with bars indicating mean ± SD. Two independent experiments were performed with a total of 12-30 fields of view. An unpaired, two-tailed, parametric student t-test with Welch’s correction was used to examine statistical differences between data sets. (**F**) Graphical representation of key findings in this study. Created with BioRender.com

## DISCUSSION

The mechanism of OC dissemination is still poorly understood; it involves the localized spreading of OC cell clusters that grow outward and detach from the primary tumor and disseminate within the peritoneal cavity. To investigate the conditions that prompt the formation of these OC outgrowths, we employed immunohistochemical stains to evaluate tissue samples representing human and PDX models of OC tumors. Immune and histochemical examination of laminin γ1 and collagens revealed that OC outgrowths were identifiable with a distinct ECM microenvironment. The ECM microenvironment observed among carcinoma cells contained abundant laminin γ1 along with lower levels of collagens expression. In contrast, the underlying extra-tumoral stroma exhibited lower laminin γ1 levels but contained dense collagen fibers. We also observed that OC outgrowths were primarily clustered away from the underlying extra-tumoral stroma, towards what we presumed was a less constrictive microenvironment. These results prompted us to use laminin- and collagen-reconstituted 3D tissue culture approaches to discern the conditions promoting the formation of OC outgrowths.

Laminin- and collagen-rich MG^[37]^ reconstitution of 2D and 3D OC cell cultures was used in combination with live-cell imaging, SEM, OCT, and rheology to examine the contribution of ECM to OC outgrowth dynamics. Live-cell imaging demonstrated that ECM reconstitution of OC spheroids, representing various OC subtypes, promoted the formation of outgrowths that, upon detachment, exhibited the ability to intercalate into mesothelial cell layers. We observed outgrowth formation in various genetically distinct OC cell lines including FNE-m-p53. ECM reconstitution did not promote the formation of outgrowths in normal, non-transformed FNE cultures, indicating that outgrowth formation is associated with transformed phenotypes of the cells of origin. OC outgrowths were observed to be initiated by cell assemblies that undergo outward translocation^[1]^ from the main spheroid. Increasing ECM concentrations led to increased elasticity of OC cell culture media, suppressed proliferation, decreased directional cell translocation, and suppressed outgrowths. Based on these results we propose a model in which OC cells favor a low elasticity ECM microenvironment to divide, translocate and form outgrowths with the potential to extend into the peritoneal cavity (**Figure 7F**). Our findings could provide reasoning for why some genetically distinct OC subtypes are superficially invasive and prefer to grow away from the ECM-dense stroma and tend to form outgrowth protrusions extending into a more fluid environment such as peritoneal ascites.

Recent IHC evaluation of OC within the fallopian tube revealed enrichment of laminin γ1 chain,^[22]^ a component of the heterotrimeric laminin αβγ molecule.^[44]^ Consistent with this study, we found that tumor cells representing OC outgrowths protruding from the ovary and omental metastases were enriched in laminin γ1. In contrast to tumor cells, the adjacent stroma displayed low levels of laminin γ1 deposition, highlighting an involvement of γ1-chain containing laminins in the regulation of cell-cell and cell-ECM^[2]^ adhesion among tumor cell assemblies competent in forming outgrowths. Laminins promote the formation of the basement membranes that support epithelial cell proliferation and collective cell migration during tissue development.^[45, 46]^ We speculate that formation of a sufficient number of cell-ECM adhesions among OC cells could contribute to tumor expansion and cell translocation, leading to the initiation of the observed outgrowths. In concurrence, our ECM-reconstituted OC spheroid model demonstrated that low concentrations of ECM were conducive in the promotion of cellular proliferation, directional cell translocation and outgrowth formation, while increased ECM concentrations were instead suppressive of the listed phenotypes. We speculate that the latter phenomenon may be attributed to increasing cell-ECM adhesion beyond the optimal point required for the processes.

Combination of SEM and OCT imaging revealed that elevation of ECM concentration, in culture, led to the formation of more thick and spaced-out protein fibers. These structural changes in ECM networks were associated with suppression of OC cell proliferation and directional translocation. These results are consistent with previous studies demonstrating collagen-fiber localization beneath OC outgrowths originating from the fallopian tube^[23]^ and within the stroma surrounding metastatic omental implants.^[47]^ Collagen fiber formation occurs when collagen concentration increases,^[48]^ indicating that extra-tumoral fibrils reflect the presence of highly elastic ECM. In addition to negative regulation of cell motion by elevation of collagen fiber content and fiber alignment,^[49]^ increasing diameter of collagen fibers^[50]^ and spacing between collagen networks^[51]^ has been shown to suppress cell motility. These data suggest that not only ECM concentration^[51]^ but also architecture of ECM is important in the regulation of cell motion.

Metastatic OC tumor cells superficially invade the mesothelial cell layer of the peritoneal membrane,^[6]^ and we have consistently observed that OC tumors growing within mouse peritoneum colonized surfaces of the omentum or small intestine and formed superficial outgrowths extending into the peritoneal cavity, with minimal invasion of the underlying stroma. Combined with our observations of outgrowth formation under low- but not high- ECM conditions, we suggest that the dynamic evolution of OC outgrowths is supported by low ECM concentration microenvironments (**Figure 3**). OC cell proliferation, translocation, and outgrowth formation were more robust under 2% MG culture conditions, indicating a possibility that elevated levels of ECM surrounding OC spheroids could provide a physical barrier that suppresses the formation of outgrowths and concomitantly hinders dissemination of OC tumor cells into the peritoneal cavity.

Recent studies have demonstrated that increasing the concentration of collagen managed to restrict single-cell translocation from cell clusters, as measured by the mean square displacement of the tracked trajectories.^[52]^ Additionally, targeting fibroblasts that produce collagens resulted in reduced collagen deposition and increased OC cluster dispersal.^[47]^ Furthermore, 2D studies of the effects of ECM concentration on cell motility have pointed to the existence of optimal ECM concentrations that support efficient directional cell migration.^[53]^ Concentrations below or above optimal levels did not stimulate fibroblast migration to the same extent.^[53]^ In agreement, our study supports this correlation by demonstrating that OC cells reconstituted with a higher concentration of ECM show a reduction in both directional cell translocation and outgrowth formation. This could be potentially explained by optimal integrin-complex turnover rates under appropriate ECM concentrations. Previously, it has been shown that high concentrations of ECM can lead to strong adhesion,^[53]^ as well as low adhesion turnover.^[54]^ Therefore, we speculate that an OC microenvironment with low ECM concentration supports the formation of adhesion complexes that are more dynamic and, thus, capable of promoting directional cell translocation. In contrast, the strong attachment of OC cells for instance due to presence of thick ECM fibers would lead to static adhesion that could impede directional cell translocation. This speculation is supported by the observation that embedding cell monolayers in high ECM increased the number of paxillin-rich focal adhesions (**S. Figure 6 and 7**).

Varying ECM concentrations were observed to modulate viscoelastic characteristics of the microenvironment and could possibly affect cell behavior,^[55]^ including the acquisition of transformed phenotypes in normal cells.^[56]^ Using rheometric analysis of our ECM- reconstituted culture media, we demonstrated that the addition of 2% MG resulted in lower elasticity as compared to the addition of 25% MG, which led to the development of thick fibrous networks and the highly elastic properties of the ECM reconstituted culture media, which is characteristic of a more solid environment (**Figure 6**). Our data demonstrated that ECM concentrations corresponding to lower elastic properties were conducive to cell proliferation, directional cell translocation, and formation of outgrowths. Thus, it may be conceivable to propose that ECM deposited on the surface of primary OC tumor cells creates a low-elasticity ECM microenvironment that provides optimal adhesion and traction force to support cell proliferation and motility of OC tumor cells. In contrast, increasing the deposition of ECM within the OC tumor microenvironment would increase adhesion and suppress cell proliferation, cell translocation, and outgrowth formation (**Figure 7F**). Our model is consistent with clinical observations that OC peritoneal dissemination is associated with tumor outgrowths that protrude from the tumor into the peritoneal cavity^[4, 22]^ through ascites ^[7, 57]^, which likely represent a low-elasticity microenvironment;^[7, 55]^ this is supported by measurements of shear moduli values of ascites, resembling those of 2% MG cultures, as well as SEM and IHC images showing fibrous networks overlaying OC cells (**S. Figure 5 and Figure 6E-I**).

OC outgrowths continue to evolve after chemotherapy because detached OC cell clusters are present within the ascitic fluid and peritoneal cavity of patients with recurrent disease.^[7]^ OC cells that have recovered from chemotherapy may be in direct contact with ECM deposited within tumors and extra-tumoral space (**Figure 2B-E**). Thus, the ECM-reconstitution model of OC outgrowths offers a clinically relevant approach to examining the dynamic processes of disease progression and recurrence. Together with the implementation of live-cell imaging, SEM, OCT, and rheology, this model can provide important insights into the role of physical characteristics of certain ECM components of OC microenvironments and their contribution to OC outgrowth formation. The presented model of OC outgrowths is limited because it does not recapitulate the full spectrum of ECM molecules associated with OC microenvironment.^[58]^ Future studies that quantitatively assess the ECM protein content *in vivo* would be necessary to guide reconstruction of ECM microenvironment for further examination of ECM contribution to OC growth.

## METHODS

### Cell Culture

Hey-A8 (obtained from Dr. Sumegha Mitra’s laboratory, University of Indiana) and Tyk-nu (obtained from Dr. Joan Brugge’s laboratory, Harvard Medical School) cells were cultured in a 1:1 ratio of Gibco™ Medium 199 (Gibco™) and MCDB105 (Sigma-Aldrich^®^) supplemented with 5% heat-inactivated fetal bovine serum (HI-FBS; Gibco™), 1% (v/v) Penicillin- Streptomycin (Gibco™) and 50 μg/ml plasmocin prophylactic (InvivoGen^®^). FNE cells (obtained from Dr. Tan Ince’s laboratory, Weill Cornell Medicine, New York) were cultured in a 1:1 ratio of Medium 199 (HiMedia^®^) and DMEM/F12 (HiMedia^®^), 3% HI-FBS (Corning^®^), 1% (v/v) Penicillin-Streptomycin (Sigma-Aldrich^®^), 0.5 ng/ml of 17 beta-estradiol (US Biological^®^), 0.2 pg/ml of triiodothyronine (Sigma-Aldrich^®^), 0.025 μg/ml all-trans retinoic acid (Beantown Chemical^®^), 20 μg/ml of insulin (Sigma-Aldrich^®^), 0.5 ng/ml of EGF (Peprotech^®^), 0.5 μg/ml hydrocortisone (Sigma-Aldrich^®^), 25 ng/ml of Cholera Toxin (Calbiochem) and 50μg/ml plasmocin prophylactic (InvivoGen^®^). ZT lung mesothelial cells (obtained from Dr. Tan Ince’s laboratory, Weill Cornell Medicine, New York) were from a benign pleural effusion. These cells were immortalized by ectopic expression of the SV40 T antigen and overexpression of human telomerase, which was fused to GFP.^[59]^Cell cultures were tested for the presence of mycoplasma every 3 months using the Uphoff and Drexler detection method.^[60]^

### IHC

Formalin-fixed, paraffin-embedded (FFPE) patient-derived xenograft tissues (PDX) were a kind gift from Dr. Ronny Drapkin, University of Pennsylvania. Sections were deparaffinized in two xylene changes for 10 minutes each, then hydrated in a graded series of ethanol (100% (v/v), 90% (v/v), 70% (v/v), 50% (v/v)) for 5 minutes each, and finally washed in ultrapure water for 5 minutes. The ABC kit, Vectastain Elite (Vector Laboratories) was used, per the manufacturer’s recommendation. The primary antibodies used were anti-PAX8 (1:1000; #10336-1-AP; Protein tech.), anti-LAMC1 (1:500; #HPA 001909; Sigma Aldrich). A peroxidase substrate (ImmPACT DAB; #SK-4105; Vector Laboratories) was used to develop and visualize staining under the microscope (Nikon ECLIPSE Ci-L equipped with a Nikon DS- Fi3 camera). The slides were rinsed in water and dehydrated in a series of ethanol solutions of increasing concentrations until 100% (v/v) ethanol was reached. The slides were then cleared in xylene, mounted with a non-aqueous mounting medium (#H-5000; Vector Laboratories), sealed using clear nail polish, and left to dry before imaging. Collagen staining was performed using a Picrosirius red staining kit (#24901; Polysciences, Inc.) per the manufacturer’s recommendation. Deparaffinization and hydration were followed by 1-hour staining in Picrosirius Red, washing in hydrochloric acid, dehydration, clearing in xylene, and mounting as described above.

### IHC Image Acquisition

Stained slides were scanned using an Olympus IX83 microscope equipped with a DP80 color camera and CellSens software. Based on scanned slides, ROIs were identified and recaptured using a Nikon Eclipse equipped with a DS-Fi3 color camera and NIS Elements D software. Images were saved in tag-image file format TIFF for further processing.

### IHC Image Processing

As previously described,^[61]^ RGB images were converted, using a weighted RGB conversion option in FIJI (Fiji is just ImageJ) software,^[62]^ to an 8-bit grayscale range representing values between black (0) and white (255). To clean out the white noise from the background, images were inverted and multiple ROIs representing tumor, stroma, or areas without tissue were selected. Mean gray values (between 0-255) were calculated for each ROI and subsequently plotted after subtracting the background signal of area without tissue. Statistical analysis was performed using the Ordinary One-way ANOVA function of GraphPad Prism [version 9.1.0 for Windows, GraphPad Software, San Diego, California USA, www.graphpad.com].

### Pathology Scoring of Laminin γ1 Expression in Human Tumors

Intensity profiles of IHC were qualitatively scored as strong (+3), moderate (+2), weak (+1), and negative (0). IHC results were recorded by a pathologist using H-scores, which were calculated by the following formula: H-score = [(0 x % negative cells) + (1 x %weakly positive cells) + (2 x %moderately positive cells) + (3 x %strongly positive cells). Data are represented as dot plots of H-scores for matched biopsy. Since the data falls into a category of a non-normal distribution, a (nonparametric) Wilcoxon rank-sum test was used to calculate significant differences between pre- and post-therapy specimens.

### ECM Reconstitution

One hundred cells per well were seeded on ultra-low attachment 96-well plates (Corning^®^), immediately centrifuged at 900 RPM for 3 minutes, and allowed to incubate until the following day. On the following day, on ice and using prechilled pipette tips, 100 μL of 4% (v/v), or 50% (v/v) Matrigel® (MG) (Corning^®^) was added to each well containing 100 μL of clustered spheroid, with a final culture volume of approximately 200 μL and final MG concentration of approximately 2% (v/v) or 25% (v/v). Matrigel lot numbers; 9294001; 9028255; 1244002; 1362001; 0288002 were used. These lot numbers represent catalogue number 354230.

### Live-Cell Imaging of 3D Cultures

Ultra-low adhesion 96-well plates containing ECM reconstituted cell cultures were placed within an Agilent^®^ BioTek^®^ LionHeart™ FX long-term imaging chamber equipped with enclosed optics, temperature, and gas exchange controls. Individual spheroids were imaged for up to 7 days with images being captured at indicated time intervals. Multiple XYZ planes of spheroids were acquired simultaneously.

### Quantification of Outgrowth Protrusion Frequency

To quantify the frequency of outgrowth protrusions by spheroids representing various FNE and OC cell lines, different laboratory members set up at least thirty spheroid cultures reconstituted with 2% (v/v) MG were prepared for each experiment. Outgrowths were subjectively defined as distinct cell populations that appeared outside of the main spheroid. Scoring was performed using a Nikon 2000 tissue culture microscope with a 10X objective.

### Mesothelial Clearance Assay

Mesothelial cells were plated on glass-bottom dishes (Mat-TEK Corporation), which had been coated with 5 μg/mL of fibronectin (Sigma-Aldrich®). GFP-expressing ZT cells were maintained in culture to form a confluent monolayer (up to 24 hours after plating). Spheroids generated with Hey-A8 were cultured for a period of seven days. Using a dissecting microscope and surgical scalpel, outgrowths were mechanically detached from their main spheroid and subsequently transferred to the co-culture containing the mesothelial monolayers. In the co-culture experiments, spheroids and their respective outgrowths were allowed to interact with a confluent mesothelial monolayer expressing green fluorescent protein (GFP). Co-culture was imaged at 1-hour intervals for up to 72 hours using an Agilent® Biotek® LionHeart™ FX (BioTek) inverted Motorized Widefield Fluorescence Microscope. Mesothelial clearance was quantified as previously described.^[39]^ The non-fluorescent area in the GFP images of the last time point of the assay, created by the penetrating cell cluster into the GFP mesothelial monolayer, was measured using FIJI ^[62]^ software. The non-fluorescent area of the final timepoint was then normalized to the area of the spheroid at the initial timepoint from the bright field channel acquisition. Data distribution and statistical analysis were conducted using GraphPad Prism [version 9.1.0 for Windows, GraphPad Software, San Diego, California USA, www.graphpad.com].

### Cell Viability and PI Incorporation Assay

The viability of both spheroid and outgrowth were quantified with a propidium iodide (PI) incorporation assay. PI is a red fluorescent dye that intercalates into double-stranded DNA but is only permeable through the compromised plasma membranes of dying/dead cells. Hey-A8 spheroids were cultured in 2% (v/v) MG and allowed to incubate for a period of 7 days. Hey- A8 spheroids were stained with PI to a final concentration of 2 µg/mL, then allowed to incubate for 30 minutes in dark. Hey-A8 spheroids were subsequently imaged with an Agilent® BioTek® LionHeart™ FX automated microscope. Z-projected images were captured with a 4X objective in both bright field and PI (590 γ excitation spectra – Texas red) channels. Following image acquisition, Z-projections for each spheroid were opened in FIJI^[62]^ and subjected to the following: Z-projections were stacked, regions of interest (ROIs) for each bright field acquisition were manually selected for both the spheroid and its associated outgrowth and selected ROIs were then superimposed over the paired PI acquisition of the same spheroid. From each Texas red channel acquisition, the mean intensity of PI was measured for both ROIs (spheroid and outgrowth) and normalized to the area of the respective ROI, thereby yielding PI mean intensity for the area of the spheroid, and for the area of the outgrowth. PI mean intensity were plotted, and statistical analysis was conducted using GraphPad Prism [version 9.1.0 for Windows, GraphPad Software, San Diego, California USA, www.graphpad.com].

### Determination of Outgrowth Length and Cellular Structure Area

GEN5 image analysis software (BioTek) was used to quantify outgrowth length and ImageJ was employed to analyze spheroid size. Before quantification, a series of bright field images, representing multiple XYZ planes of a discrete cellular structure, was collapsed to generate maximal projections of a spheroid and its protruding outgrowth. Outgrowth protrusion was identified as a structure that extended beyond the main spheroid body, and a line was drawn from the tip across the longer axis of the outgrowth, terminating at the junction between outgrowth protrusion and the spheroid. The size of the cellular structure was calculated using an in-house ImageJ macro. Binary masks were created to separate a single cellular structure from the non-cellular background.

### Cryo-SEM Sample Preparation and Imaging

All samples for cryo-scanning electron microscopy (SEM) were prepared by using a Leica EM HPM100 high-pressure freezing (HPF) system. The HPF planchettes were washed in ethanol and exposed to an oxygen plasma for ten minutes before use. The frozen-hydrated samples were stored in liquid nitrogen. A Leica VCT-100 system was used for subsequent cryo-transfer and cryo-imaging. Samples were transferred and coated with sputtered gold (Au) (2.5 nm) under cryogenic conditions (T < -135 °C) by a Leica EM MED020 system. Before SEM imaging, sublimation was used to create topographic contrast by slightly warming frozen-hydrated samples. SEM imaging was done by using a Zeiss Auriga Crossbeam FIB-SEM equipped with a Schottky field-emission electron gun (FEG) and an Oxford Max-80 ultrathin window (UTW) silicon-drift detector (SDD) interfaced to an Oxford INCA EDS system. Secondary electron imaging (Everhart-Thornley detector) was done by using 2 keV electrons and a 2.5 nm Au coating.

### Assessment of ECM network in MG with OCT

A spectral-domain OCT system with a central wavelength of approximately 850 mm was used for MG imaging. The system provides an axial resolution of approximately 9 μm in biological samples (1.4 refractive indexes assumed) and a transverse resolution of approximately 5 μm. Parameters for imaging of 2% (v/v) and 25% (v/v) MG samples were kept identical. Data processing focused on the characterization of the spatial frequency over depth, similar to a previously developed method.^[63]^ Briefly, with the dB intensity A-scan and the identified sample surface, the intensity profile over 0.4 mm starting at approximately 26 μm below the sample surface was utilized for analysis. The slope of the signal that represents the optical attenuation over depth was removed. Through a fast Fourier transform, the amplitude spectrum of the spatial frequency was obtained, and an exponential fit of the local peaks was performed. The exponential decay coefficient was calculated as the measure of the spatial frequency over depth and was used to compare the 2% (v/v) and 25% (v/v) MG samples. A higher decay coefficient represents relatively stronger low-frequency components of the spatial frequency spectrum.

### Rotational Rheology

The linear viscoelastic material functions, the storage modulus, G’, and the loss modulus, G”, of the 2% (v/v), 25% (v/v) MG samples, as well as ascitic fluids isolated from OC patients, were characterized using an Advanced Rheometric Expansion System (ARES) rheometer available from TA Instruments of New Castle, DE. The rotational rheometer was used with stainless steel parallel disks with a 25 mm diameter and had a force rebalance transducer 0.2K- FRTN1. The actuator of the ARES is a DC servomotor with a shaft supported by an air bearing with an angular displacement range of 5×10^−6^ to 0.5 rad, and an angular frequency range of 1×10^−5^ to 100 rad/s. The angular velocity range is 1×10^−6^ to 200 rad/s. The sample loading procedure was the same for all the experiments, and the gap height between two disks was kept constant at 1 mm. A sufficient volume of sample was used to fill the gap between parallel disks, and the linear viscoelastic properties of the samples were collected as a function of frequency at a constant strain of 50% and room temperature. The samples were not pre-sheared. For discussion of dynamic property characterization of complex fluids see, for example, Bird, R.B., Armstrong, R.C., and Hassager, O. “Dynamics of Polymeric Liquids. Wiley, 1987.”

### Matrigel® (MG) Overlay

MG was diluted with an appropriate prechilled medium and a sterile and precooled pipette tip to a final concentration of 2% (v/v) or 25% (v/v). After cell attachment, the medium was aspirated from the 96-well plates and the medium/MG mixture was overlaid onto the cell monolayer. MG-treated cultures were allowed to incubate overnight before proceeding with live-cell time-lapse imagining or incubated for 72 h before fixation and IF staining. The procedure as described produces a coat of matrix that gels at 37°C and attaches to the upper surface of the monolayer.^[41]^

### Processing of spheroids for EdU labeling

Hey-A8 spheroids were fixed in a 4% (v/v) paraformaldehyde solution (Sigma-Aldrich^®^) and dehydrated in ethanol. HistoGel™ (Thermo Fisher Scientific^®^) was liquified, and spheroids were mixed with HistoGel™ and left to solidify in a biopsy cassette. HistoGel™ blocks were fixed in 10% (v/v) neutral-buffered formalin overnight, then dehydrated in ethanol and cleared in xylene. Processed HistoGel™ blocks were then embedded in paraffin and stored at -20 °C until the time of sectioning. Paraffin blocks were sectioned at a thickness of 5 μm using a microtome (Leica).

### EdU incorporation assay

MG reconstituted Hey-A8 cells were incubated with 10 mM Click-iT™ EdU [Thermo Fisher Scientific^®^, catalog #: C10340] for 4 hours. Cells were fixed with 4% (v/v) PFA for 1 hour at room temperature, embedded into HistoGel™, and sectioned. Sections were permeabilized with 1X PBS containing 0.5% (v/v) Triton-X. Residual Triton-X was removed by washing twice with 1X PBS supplemented with 3% (w/v) BSA. To label the incorporated EdU, sections were incubated in the dark for 30 minutes with a Click-iT reaction cocktail prepared fresh (<15 minutes before labeling). The Click-iT reaction cocktail was then removed, and sections were washed with 1X PBS. To label total cell nuclei, sections were incubated in the dark with 10 μg/mL Hoechst 33342 in 1X PBS for 30 minutes. After washing out the Hoechst dye with 1X PBS, sections were mounted with aqueous mounting media and sealed by coverslips. The sections were imaged on ZEISS LSM-880 confocal microscope using a 10X objective. Images of EdU-labeled nuclei were acquired using far-red laser illumination (λ = 647 nm) and total nuclei were captured by violet-blue laser illumination (λ = 405 nm). Images were analyzed using the open-source software FIJI.^[62]^

### Cell proliferation, cell motion imaging, and quantifications

Monolayers of FNE-m-p53 or Hey-A8 cells expressing GFP or mKO2, respectively, were overlaid with MG. Time-lapse imaging was performed on an Agilent® BioTek® Lionheart™ FX automated microscope using a 10X objective and a maximal capture interval of 22 minutes. After the acquisition, time-lapse images were background subtracted (radius 50 pixels) using FIJI^[62]^ image analysis software. To analyze cell proliferation and movement within MG- overlaid cell cultures, we used an ImageJ plugin, TrackMate,^[64]^ to perform single-particle tracking (SPT). TrackMate offers multiple modes of analysis and based on both our cell size and the relatively slow movement of cells, we selected and applied the following parameters in TrackMate: LoG (Laplacian of Gaussian) detector, the spot diameter of 35 pixels, threshold of 10, median filter and LAP tracker with 15-pixel frame linking, 2-frame gap distance and track-segment splitting. We extracted the spot count in the desired time frames (one count every 12 hours or 33 frames) to obtain the number of cells or spots in the region of interest at those time points. We then divided the number of cells in each time point by the number of cells at the starting time point (time = 0) to normalize and plot it as a measure of fold change. An unpaired, two-tailed t-test was used to assess the significance of fold change in cell number between the first and final time points. Plotting and statistical analysis were conducted using GraphPad Prism [version 9.1.0 for Windows, GraphPad Software, San Diego, California USA, www.graphpad.com]. To quantify track displacement, we calibrated the images by converting output in pixels into micrometers, and frames into hours. For visualization of local tracks, we used 50-frame-depth-, time-color-coded tracks overlayed on a GFP or RFP channel time-lapse. For trajectory classification, we used the TraJClassifier plugin.^[65]^ The plugin uses TrackMate output data and trajectories and computes a confinement ratio representing cell directionality over time. After all the trajectories were computed into confinement ratios, we plotted the data using Prism GraphPad Prism 9.1.0. Trajectories from multiple fields of view were plotted and directed-motion trajectories were represented as confinement ratio values. Each dot in a dot plot represents one field of view with the horizontal line depicting the mean of all fields of view per condition. Statistical analysis was computed using an unpaired, two-tailed t-test.

### Ascites and solid carcinoma tissue collection

Ascitic fluids were collected from patients with ovarian cancer either by paracentesis under local anesthesia or at the very beginning of surgery. Ascites was centrifuged at 1,100 x *g* for 10 min at room temperature to remove cell clusters, then aliquoted and stored at -80 °C until analysis. All solid tissue collections were performed during surgery. All procedures performed in studies involving human participants were following the ethical standards of the institutional and national research committee and with the 1964 Helsinki declaration and its later amendments or comparable ethical standards. Patients provided a signed informed consent, approved by the Ethics Review Board of Poznań University of Medical Sciences (Consent No 737/17).

### Immunofluorescence staining

MG overlayed 2D cultures or spheroid cultures were washed thrice with PBS, fixed in 4% paraformaldehyde for 15-30 min at room temperature. Following blocking with 2.5% normal goat serum for 60 min, cells were incubated with indicated primary antibodies for 1 h at room temperature. Following PBS washes, cells were incubated with secondary antibody tagged with fluorescein diluted at 1:500 in PBS. For nuclear staining, Hoechst was used. Immunostained samples were examined under ZEISS LSM-880, the confocal laser scanning microscope (CLSM).

### N-cadherin quantification

Still images of HeyA8 spheroids reconstituted with 2% or 25% MG and stained with N-cadherin and DAPI were acquired on laser scanning confocal microscope (ZEISS LSM 880) using 20x objective. Still images were collected as Z-stacks which were later projected after acquisition using standard deviation projection mode with image analysis software Fiji.^[62]^ As the Hey-A8 spheroids were too large to fit into a single image frame using the 20x objective, we focused on imaging the spheroid edges, capturing different edges of spheroids in multiple frames. To quantify the level of N-cadherin staining strictly at the spheroid edge, we manually selected the region of the edge and measured the raw integrated density of the N-cadherin channel (547 nm) and the DAPI channel (405 nm). To account for the different size of the region and varying number of cells in each region, we first normalized the raw integrated density to the region area (in pixels) and then normalized that value from the N-cadherin channel (547 nm) to the DAPI channel (405 nm). Normalized N-cadherin value was plotted using Prism GraphPad Prism [version 9.1.0 for Windows, GraphPad Software, San Diego, California USA, www.graphpad.com]. Each value on the plot represents one spheroid edge with the bar height depicting the mean of all edges taken per condition with error bars as standard deviation. A total of 31 – 43 edges were quantified per condition, corresponding to 16 – 19 Hey-A8 spheroids per condition. Statistical analysis was performed using the Ordinary One-way ANOVA function of GraphPad Prism [version 9.1.0 for Windows, GraphPad Software, San Diego, California USA, www.graphpad.com].

### Focal adhesion quantification

Still images of FNE m-p53 and Hey-A8 cells overlayed with 2% or 25% Matrigel and stained with paxillin, phalloidin (F-actin) and Hoechst 33342 were acquired on laser scanning confocal microscope (ZEISS LSM 880) using 20x objective. Still images were collected as Z-stacks which were later projected after acquisition using maximum projection mode with image analysis software Fiji.^[62]^ To quantify number of focal adhesion foci we used an ImageJ plugin TrackMate,^[64]^ a readily available tool to perform a range of operations from particle detection to single particle tracking (SPT). TrackMate offers multiple modes of analysis and based on intensity and size of our paxillin foci, we selected and applied the following parameters in TrackMate: LoG (Laplacian of Gaussian) detector, estimated object diameter of 1 micron, quality threshold 20, with applied median filter. We applied this setting to each ROI we imaged. To account for the varying number of cells in each ROI, we normalized the total number of focal adhesion (paxillin) foci detected to the total number of nuclei in the respective ROI, which we counted using the TrackMate thresholding detector with auto intensity threshold. Normalized number of focal adhesion foci was then plotted using Prism GraphPad Prism [version 9.1.0 for Windows, GraphPad Software, San Diego, California USA, www.graphpad.com]. Each value on the plot represents one ROI with the bar height depicting the mean of all ROIs per condition, with error bars as standard deviation. A total of 10 ROIs were quantified per condition, ranging between 12 – 376 cells per ROI. Statistical analysis was performed using the Ordinary One-way ANOVA function of GraphPad Prism [version 9.1.0 for Windows, GraphPad Software, San Diego, California USA, www.graphpad.com].

### PAX8 quantification for stroma invasion

Using FIJI (Fiji is just ImageJ) software,^[62]^ multiple ROIs representing tumor, stroma, or Villi small intestine were selected. Integrated density "IntDen" (the product of Area and Mean Gray Value)^[62]^ was calculated for each ROI subsequently plotted. Statistical analysis was performed using the Ordinary One-way ANOVA function of GraphPad Prism [version 9.1.0 for Windows, GraphPad Software, San Diego, California USA, www.graphpad.com]. ANOVA was followed by Tukey post-hoc analysis.

### Statistical analysis

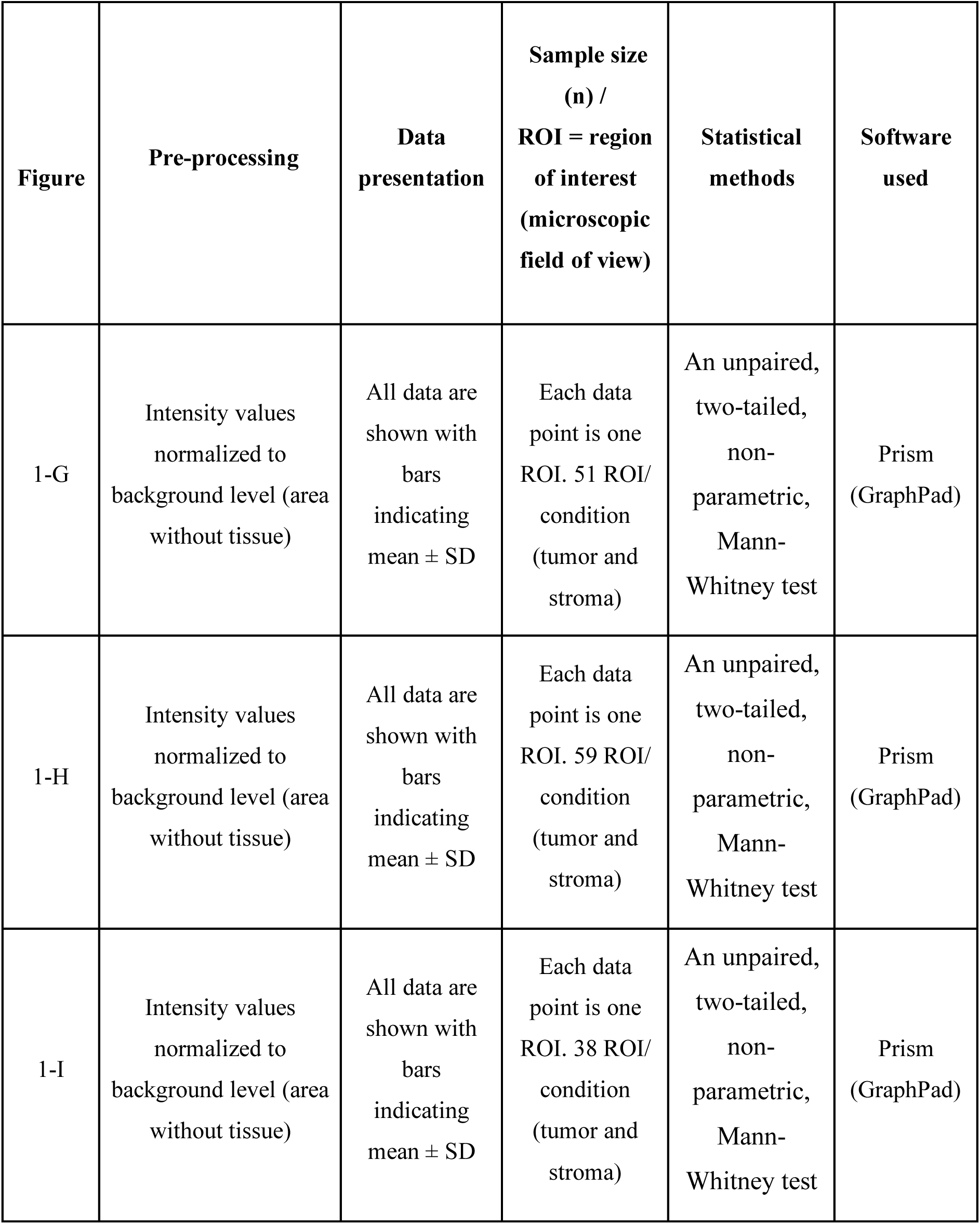
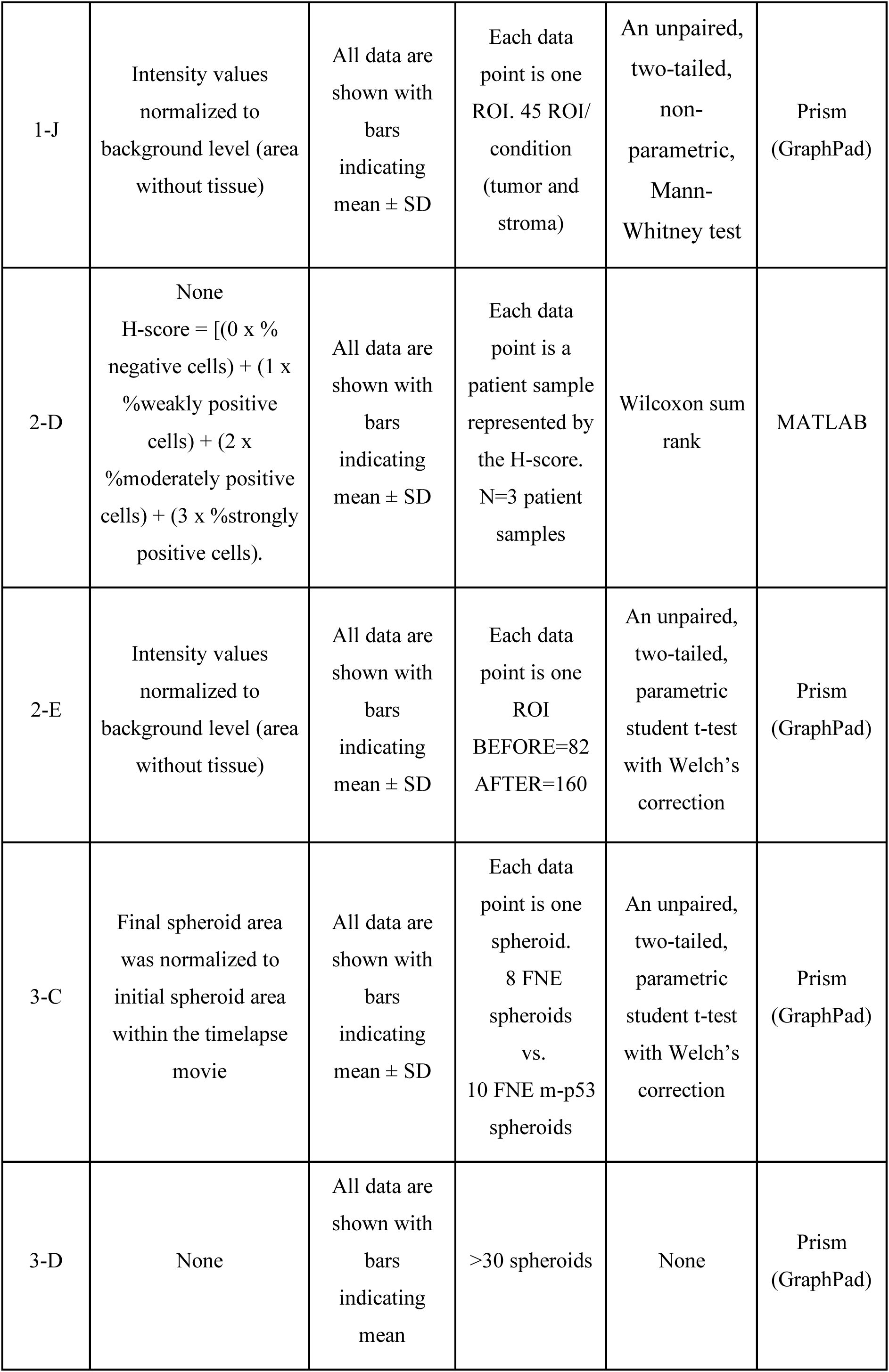
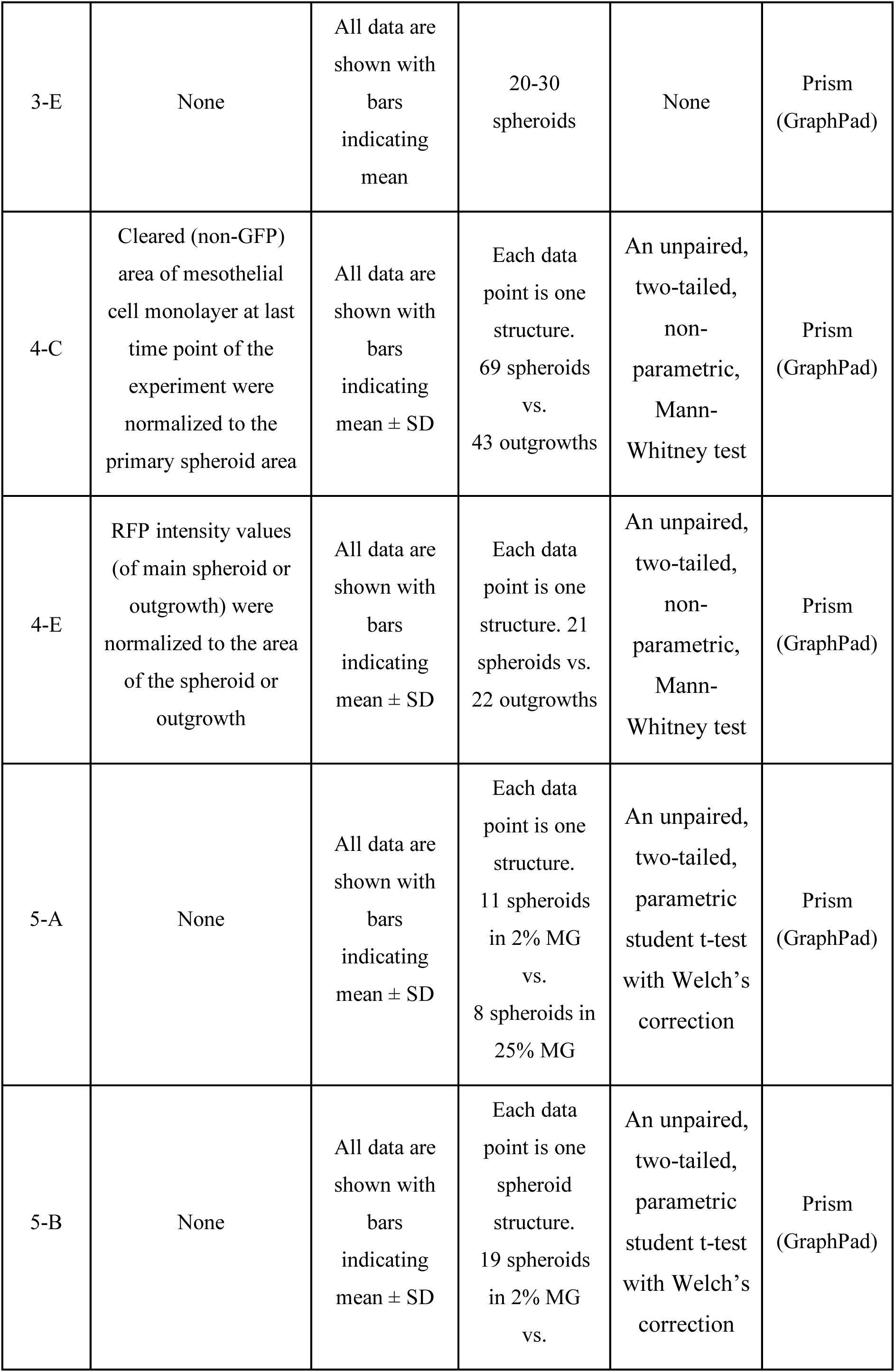
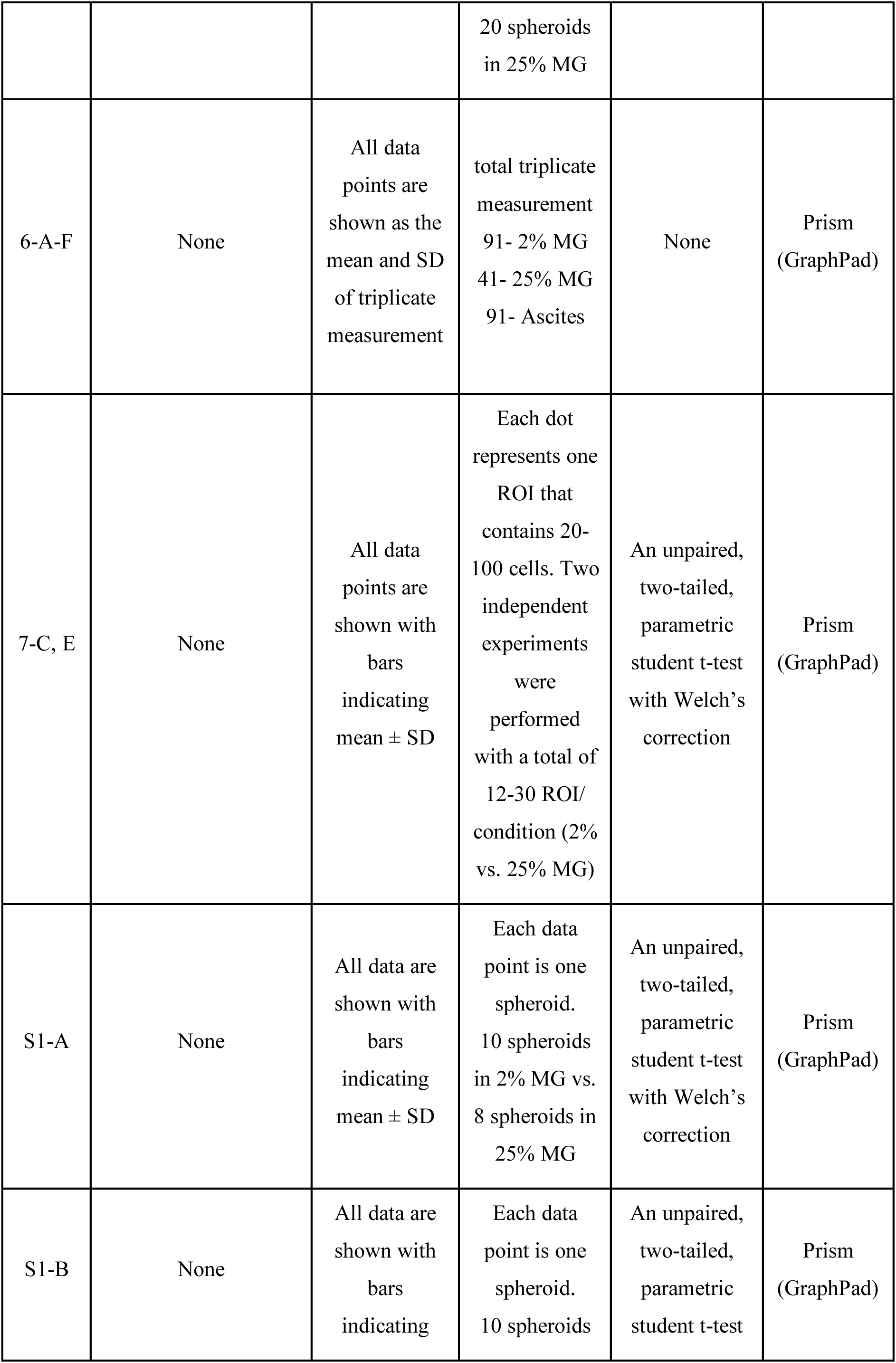
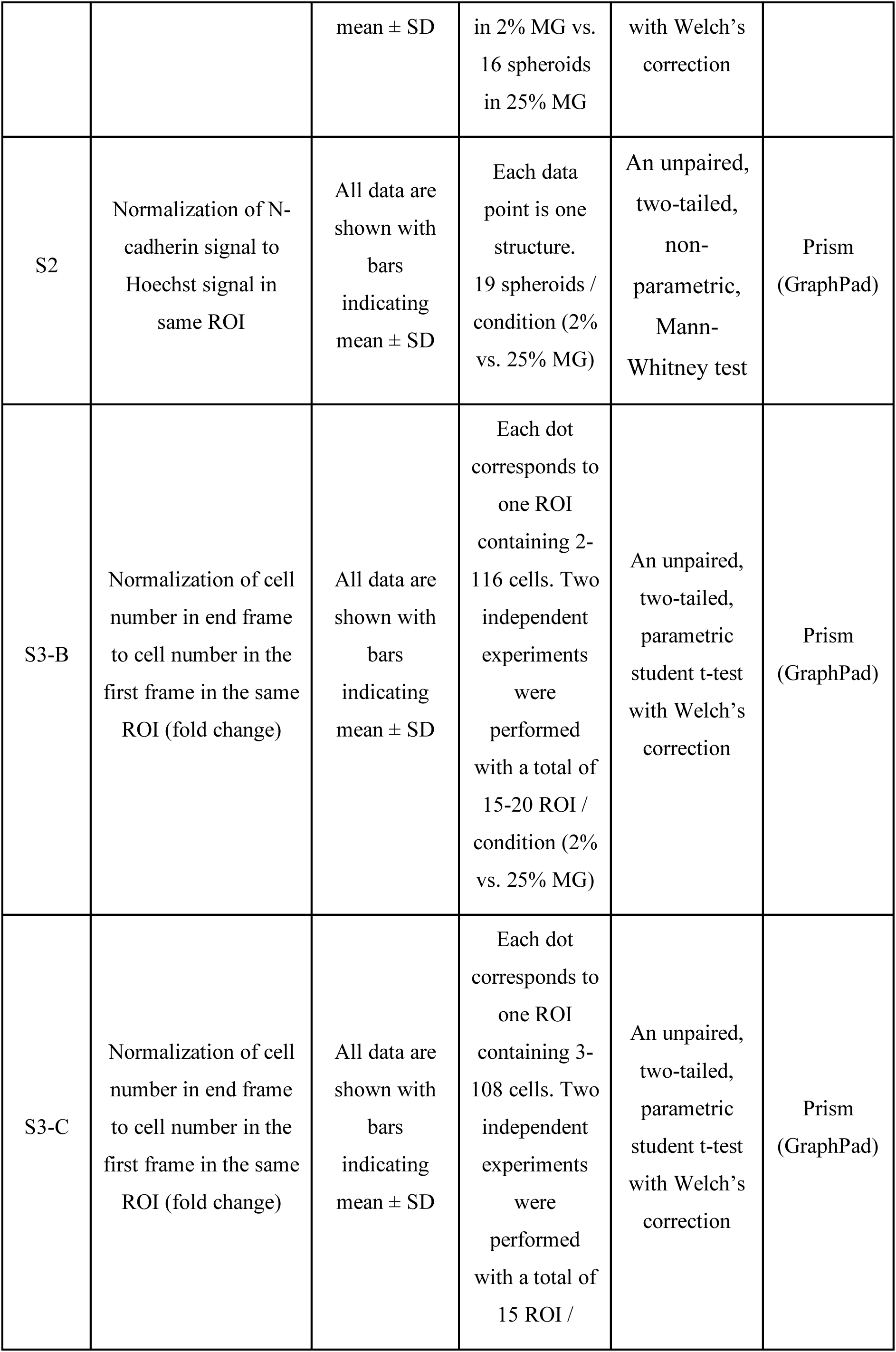
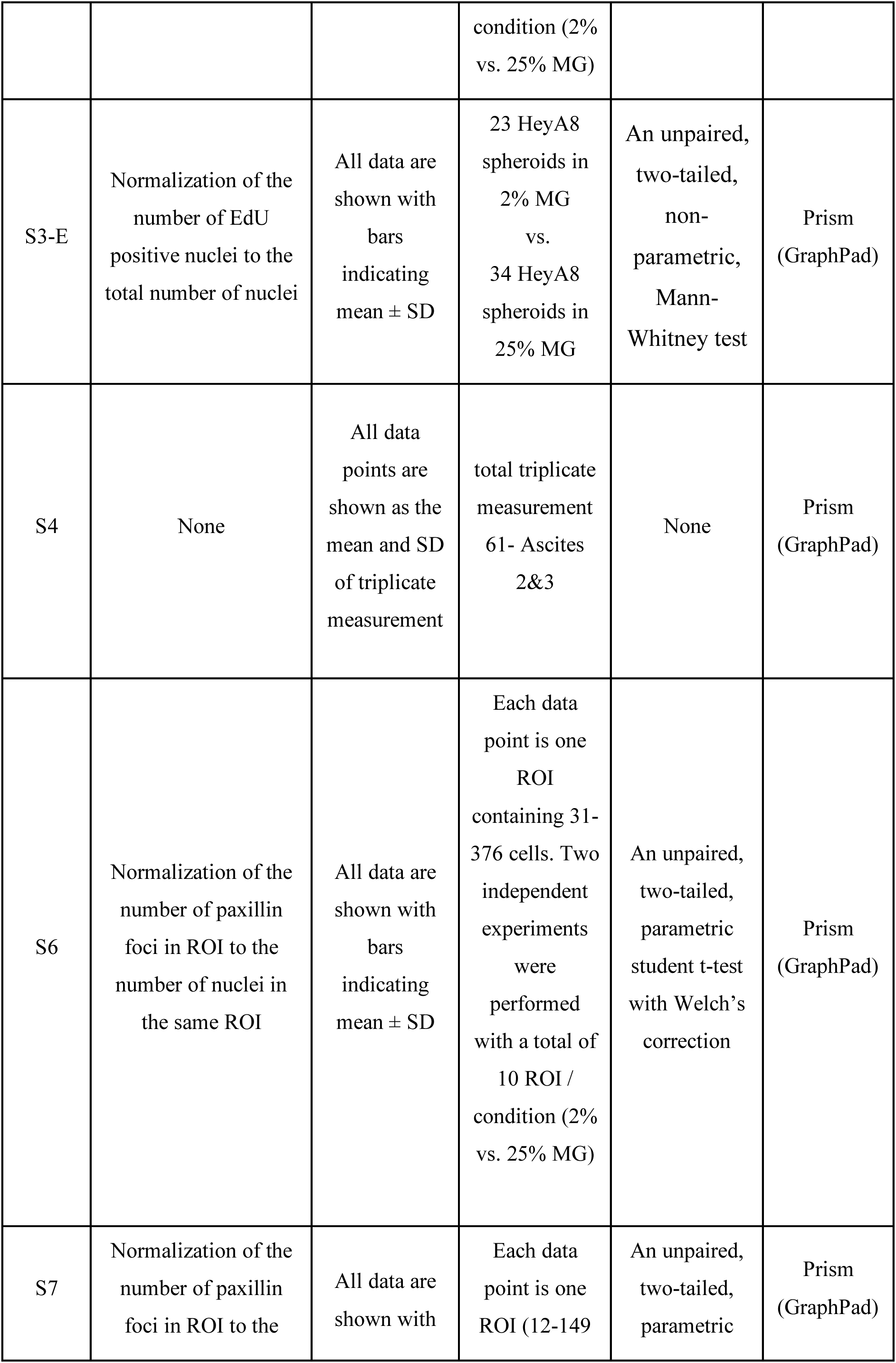
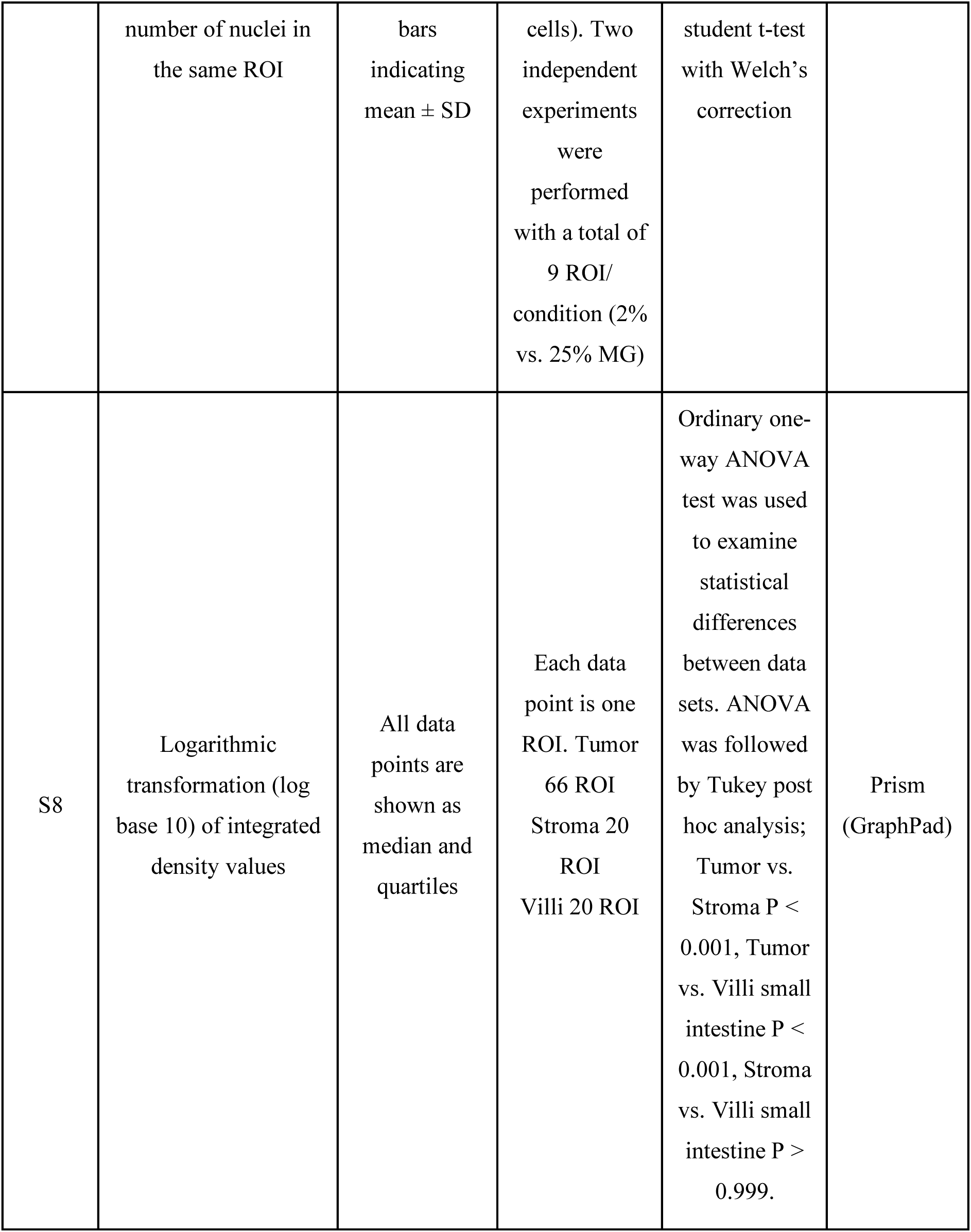

| Figure | Pre-processing | Data presentation | Sample size (n) / ROI = region of interest (microscopic field of view) | Statistical methods | Software used |
| --- | --- | --- | --- | --- | --- |
| 1-G | Intensity values normalized to background level (area without tissue) | All data are shown with bars indicating mean ± SD | Each data point is one ROI. 51 ROI/ condition (tumor and stroma) | An unpaired, two-tailed, non-parametric, Mann-Whitney test | Prism (GraphPad) |
| 1-H | Intensity values normalized to background level (area without tissue) | All data are shown with bars indicating mean ± SD | Each data point is one ROI. 59 ROI/ condition (tumor and stroma) | An unpaired, two-tailed, non-parametric, Mann-Whitney test | Prism (GraphPad) |
| 1-I | Intensity values normalized to background level (area without tissue) | All data are shown with bars indicating mean ± SD | Each data point is one ROI. 38 ROI/ condition (tumor and stroma) | An unpaired, two-tailed, non-parametric, Mann-Whitney test | Prism (GraphPad) |
| 1-J | Intensity values normalized to background level (area without tissue) | All data are shown with bars indicating mean ± SD | Each data point is one ROI. 45 ROI/condition (tumor and stroma) | An unpaired, two-tailed, non-parametric, Mann-Whitney test | Prism (GraphPad) |
| 2-D | None<br>H-score = [(0 x % negative cells) + (1 x %weakly positive cells) + (2 x %moderately positive cells) + (3 x %strongly positive cells)]. | All data are shown with bars indicating mean ± SD | Each data point is a patient sample represented by the H-score. N=3 patient samples | Wilcoxon sum rank | MATLAB |
| 2-E | Intensity values normalized to background level (area without tissue) | All data are shown with bars indicating mean ± SD | Each data point is one ROI<br>BEFORE=82<br>AFTER=160 | An unpaired, two-tailed, parametric student t-test with Welch's correction | Prism (GraphPad) |
| 3-C | Final spheroid area was normalized to initial spheroid area within the timelapse movie | All data are shown with bars indicating mean ± SD | Each data point is one spheroid.<br>8 FNE spheroids vs. 10 FNE m-p53 spheroids | An unpaired, two-tailed, parametric student t-test with Welch's correction | Prism (GraphPad) |
| 3-D | None | All data are shown with bars indicating mean | >30 spheroids | None | Prism (GraphPad) |
| 3-E | None | All data are shown with bars indicating mean | 20-30 spheroids | None | Prism (GraphPad) |
| 4-C | Cleared (non-GFP) area of mesothelial cell monolayer at last time point of the experiment were normalized to the primary spheroid area | All data are shown with bars indicating mean ± SD | Each data point is one structure.<br>69 spheroids vs.<br>43 outgrowths | An unpaired, two-tailed, non-parametric, Mann-Whitney test | Prism (GraphPad) |
| 4-E | RFP intensity values (of main spheroid or outgrowth) were normalized to the area of the spheroid or outgrowth | All data are shown with bars indicating mean ± SD | Each data point is one structure. 21 spheroids vs. 22 outgrowths | An unpaired, two-tailed, non-parametric, Mann-Whitney test | Prism (GraphPad) |
| 5-A | None | All data are shown with bars indicating mean ± SD | Each data point is one structure.<br>11 spheroids in 2% MG vs.<br>8 spheroids in 25% MG | An unpaired, two-tailed, parametric student t-test with Welch's correction | Prism (GraphPad) |
| 5-B | None | All data are shown with bars indicating mean ± SD | Each data point is one spheroid structure.<br>19 spheroids in 2% MG vs. | An unpaired, two-tailed, parametric student t-test with Welch's correction | Prism (GraphPad) |
|  |  |  | 20 spheroids<br>in 25% MG |  |  |
| 6-A-F | None | All data<br>points are<br>shown as the<br>mean and SD<br>of triplicate<br>measurement | total triplicate<br>measurement<br>91- 2% MG<br>41- 25% MG<br>91- Ascites | None | Prism<br>(GraphPad) |
| 7-C, E | None | All data<br>points are<br>shown with<br>bars<br>indicating<br>mean ± SD | Each dot<br>represents one<br>ROI that<br>contains 20-100 cells. Two<br>independent<br>experiments<br>were<br>performed<br>with a total of<br>12-30 ROI/<br>condition (2%<br>vs. 25% MG) | An unpaired,<br>two-tailed,<br>parametric<br>student t-test<br>with Welch's<br>correction | Prism<br>(GraphPad) |
| S1-A | None | All data are<br>shown with<br>bars<br>indicating<br>mean ± SD | Each data<br>point is one<br>spheroid.<br>10 spheroids<br>in 2% MG vs.<br>8 spheroids in<br>25% MG | An unpaired,<br>two-tailed,<br>parametric<br>student t-test<br>with Welch's<br>correction | Prism<br>(GraphPad) |
| S1-B | None | All data are<br>shown with<br>bars<br>indicating | Each data<br>point is one<br>spheroid.<br>10 spheroids | An unpaired,<br>two-tailed,<br>parametric<br>student t-test | Prism<br>(GraphPad) |
| | | mean ± SD | in 2% MG vs.<br>16 spheroids<br>in 25% MG | with Welch's<br>correction | |
| S2 | Normalization of N-cadherin signal to Hoechst signal in same ROI | All data are shown with bars indicating mean ± SD | Each data point is one structure.<br>19 spheroids / condition (2% vs. 25% MG) | An unpaired, two-tailed, non-parametric, Mann-Whitney test | Prism (GraphPad) |
| S3-B | Normalization of cell number in end frame to cell number in the first frame in the same ROI (fold change) | All data are shown with bars indicating mean ± SD | Each dot corresponds to one ROI containing 2-116 cells. Two independent experiments were performed with a total of 15-20 ROI / condition (2% vs. 25% MG) | An unpaired, two-tailed, parametric student t-test with Welch's correction | Prism (GraphPad) |
| S3-C | Normalization of cell number in end frame to cell number in the first frame in the same ROI (fold change) | All data are shown with bars indicating mean ± SD | Each dot corresponds to one ROI containing 3-108 cells. Two independent experiments were performed with a total of 15 ROI / | An unpaired, two-tailed, parametric student t-test with Welch's correction | Prism (GraphPad) |
|  |  |  | condition (2% vs. 25% MG) |  |  |
| S3-E | Normalization of the number of EdU positive nuclei to the total number of nuclei | All data are shown with bars indicating mean ± SD | 23 HeyA8 spheroids in 2% MG vs. 34 HeyA8 spheroids in 25% MG | An unpaired, two-tailed, non-parametric, Mann-Whitney test | Prism (GraphPad) |
| S4 | None | All data points are shown as the mean and SD of triplicate measurement | total triplicate measurement 61- Ascites 2&3 | None | Prism (GraphPad) |
| S6 | Normalization of the number of paxillin foci in ROI to the number of nuclei in the same ROI | All data are shown with bars indicating mean ± SD | Each data point is one ROI containing 31-376 cells. Two independent experiments were performed with a total of 10 ROI / condition (2% vs. 25% MG) | An unpaired, two-tailed, parametric student t-test with Welch's correction | Prism (GraphPad) |
| S7 | Normalization of the number of paxillin foci in ROI to the | All data are shown with | Each data point is one ROI (12-149 | An unpaired, two-tailed, parametric | Prism (GraphPad) |
| | number of nuclei in the same ROI | bars indicating mean ± SD | cells). Two independent experiments were performed with a total of 9 ROI/condition (2% vs. 25% MG) | student t-test with Welch's correction | |
| S8 | Logarithmic transformation (log base 10) of integrated density values | All data points are shown as median and quartiles | Each data point is one ROI. Tumor 66 ROI Stroma 20 ROI Villi 20 ROI | Ordinary one-way ANOVA test was used to examine statistical differences between data sets. ANOVA was followed by Tukey post hoc analysis; Tumor vs. Stroma P < 0.001, Tumor vs. Villi small intestine P < 0.001, Stroma vs. Villi small intestine P > 0.999. | Prism (GraphPad) |

### Reagents and materials

**Reagents and materials:**
| Item | Cat. No. | Vendor |
| --- | --- | --- |
| PAX8 | 10336-1-AP | Protein tech. |
| LAMC1 | HPA 001909 | Sigma Aldrich |
| VECTASTAIN Elite ABC HRP | PK-6101 | vector laboratories |
| ImmPACT DAB | SK-4105 | vector laboratories |
| Hematoxylin QS | H-3404-100 | vector laboratories |
| picrosirius red | 24901 | Polysciences Inc. |
| non-aqueous mounting medium | H-5000 | vector laboratories |
| Aqueous mounting medium | H-5501 | vector laboratories |
| Medium 199 | AT014-5L | Himedia |
| MCDB105 | M6396-10X1L | Sigma Aldrich |
| Heat Inactivated Fetal bovine serum | 10438-026500mL | Gibco |
| Plasmocin prophylactic | ant-mpp | In vivo Gen |
| Penicillin-Streptomycin | 30-002-CI | Corning |
| DMEM/F12 | AT147A-10L | Himedia |
| 17 beta-estradiol | 15600 | USB |
| triiodothyronine | T6397-100MG | Sigma Aldrich |
| all-trans retinoic acid | 216220-250MG | BeanTown Chemical |
| EGF | 100-26-25UG | Peprotech |
| hydrocortisone | H088-1G | Sigma Aldrich |
| cholera toxin | C8052 | Sigma Aldrich |
| Matrigel | 354230 | Corning |
| ultra-low attachment 96-well plates | 7007 | Corning |
| Trypsin | 25-053-CI | Corning |
| Insulin | I9278-5ML | Sigma Aldrich |
| Histogel | HG-4000-012 | Thermo Scientific |
| Formalin | VT450C | Lam Chem |
| Xylene | 89370-088 | VWR |
| Ethanol | 241000200CSGL | GREENFIELD GLOBAL INC |
| PBS | 21-040-CV | Corning |
| Propidium iodide | P1304MP | Molecular Probes |
| EDU kit | C10640 | LIFE TECHNOLOGIES |
| N-Cadherin | C3865 | Sigma Aldrich |
| Paxillin | 610051 | BD Pharmingen |
| Phalloidin | A12380 | ThermoFisher |
| Alpha smooth muscle actin | ab7817 | Abcam |
| Triton-X and Tween-20 | 0777-1L | VWR |
| Alexa fluor 488 | A11029 | Invitrogen |
| Alexa fluor 546 | A11003 | Invitrogen |
| Alexa fluor 568 | A-11031 | Invitrogen |

## Author Contributions

S.A., T.P. and M.I. designed and conceptualized the studies with significant input from S.F., P.B., D.C., and D.K. IHC analysis of human tumors and PDX tissue sections were performed by S.A., and T.P. performed cell tracking experiments and image quantification. S.F. performed mesothelial clearance assays. P.B., L.Q., and D.Kh. designed and performed shear-stress rheometric analyses. P.B. and W.L. designed and performed electron microscopy experiments. M.R., P.J., E.D., E.N-M. and M.P-Z., identified human tumor samples and performed pathologic analysis of laminin γ1 expression in tumors representing disease before and after chemotherapy. S.A., T.P., and M.I. wrote a manuscript with significant input (methodology description) from S.F., P.B., L.Q., D.Kh., S.W., and M.P- Z.

## Acknowledgments

We would like to thank Dr. Shang Wang for assisting with OCT experiments and help with writing the relevant methodology and the interpretation of the results. We thank the Stevens Institute of Technology, Multiscale Imaging Center led by Dr. Matthew Libera. We are especially grateful for technical help with sample preparation and processing that was provided by Dr. Tsengming Chou. We thank Thomas Cattabiani for editorial help with the manuscript preparation. This study was supported by NIH/NCI R21CA256615 (M.I.) and Kaleidoscope of Hope Ovarian Cancer Foundation (M.I.).

## Supplementary Figures

**S. Figure 1.**
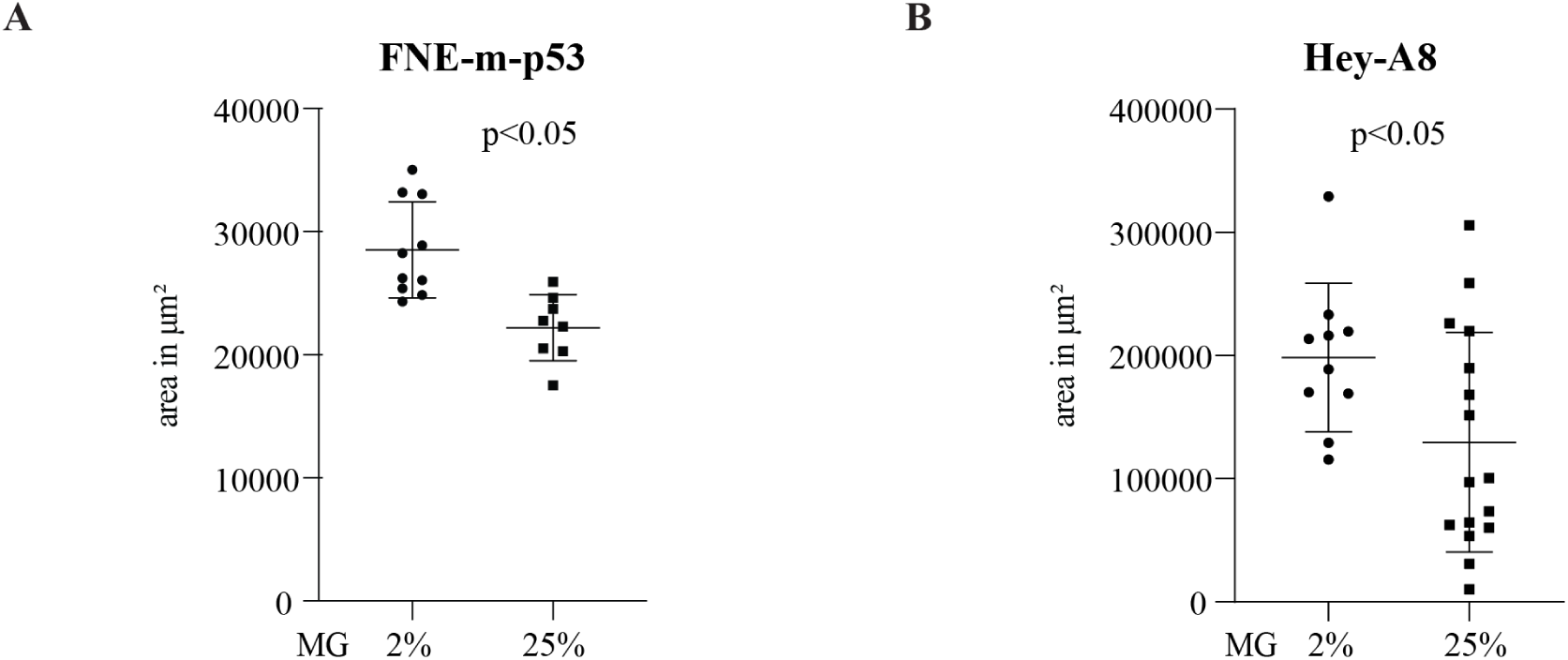
Elevation of ECM concentration suppresses FNE-m-p53 and Hey-A8 structure expansion. Quantification of an area in (**A**) FNE-m-p53 or (**B**) Hey-A8 spheroids reconstituted with 2% or 25% MG. Each dot represents a spheroid (8-16 spheroid/condition) measured 9 days after starting the culture. All data are shown with bars indicating mean ± SD. An unpaired, two-tailed, parametric student t-test with Welch’s correction was used to examine statistical differences between data sets.

**S. Figure 2.**
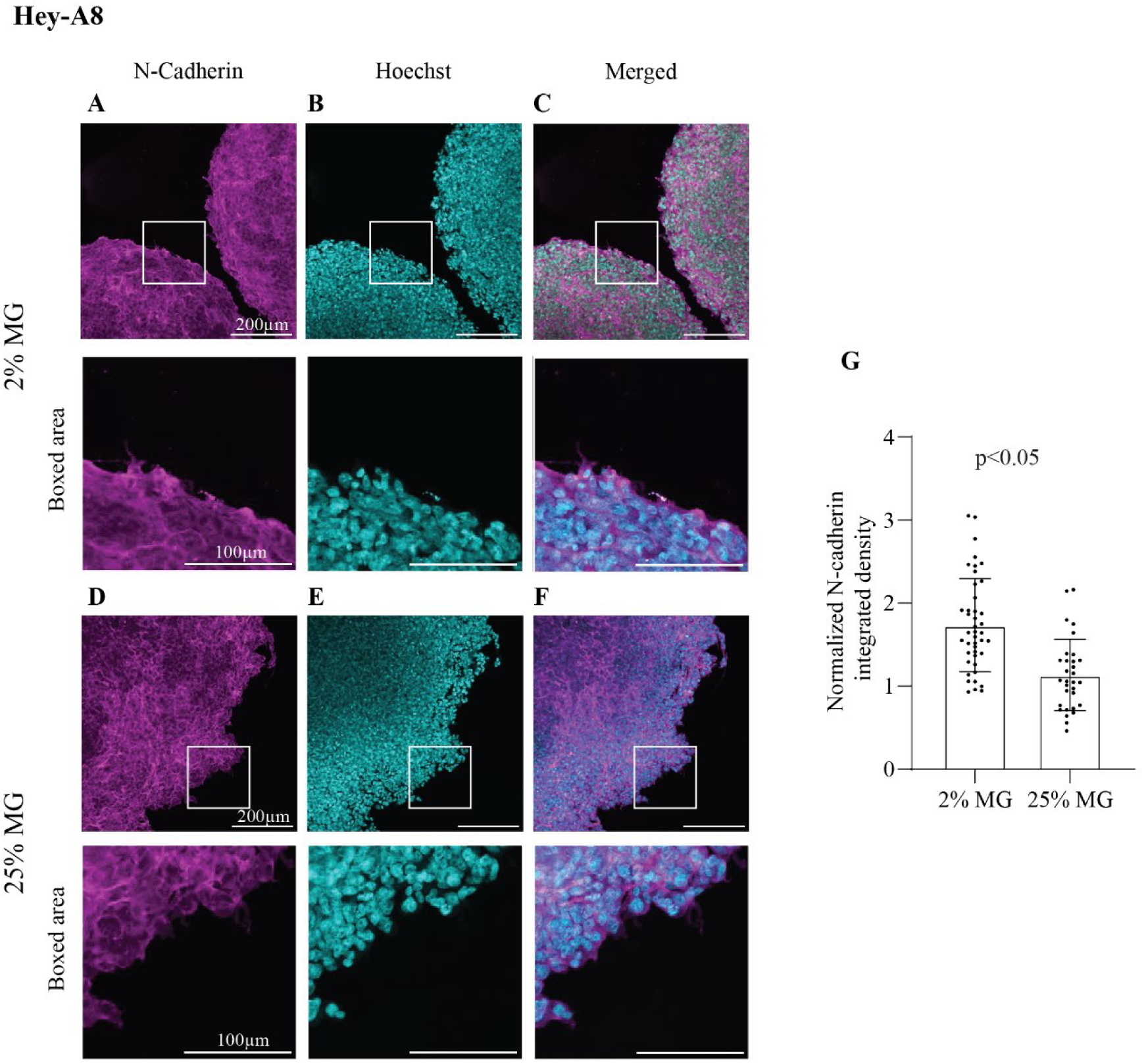
Increased ECM concentration leads to reduced N-cadherin expression at the 3D OC spheroid surface edge. Hey-A8 spheroids were reconstituted with (**A, B, C**) 2% MG or (**D, E, F**) 25% MG and stained with (**A, D**) antibodies against N-cadherin, and (**B, E**) Hoechst 33342 dye. (**G**) Quantification of N-cadherin expression on the edges of Hey-A8 spheroids reconstituted with 2% or 25% MG, respectively. Each data point is one ROI (1-5 spheroid edges). Nineteen spheroids were analyzed/ condition. All data points are shown with bars indicating mean ± SD. An unpaired, two-tailed, non-parametric, Mann-Whitney test was used to examine statistical differences between data sets. Integrated density values of N-cadherin were normalized to integrated density values of corresponding Hoechst 33342 signals.

**S. Figure 3.**
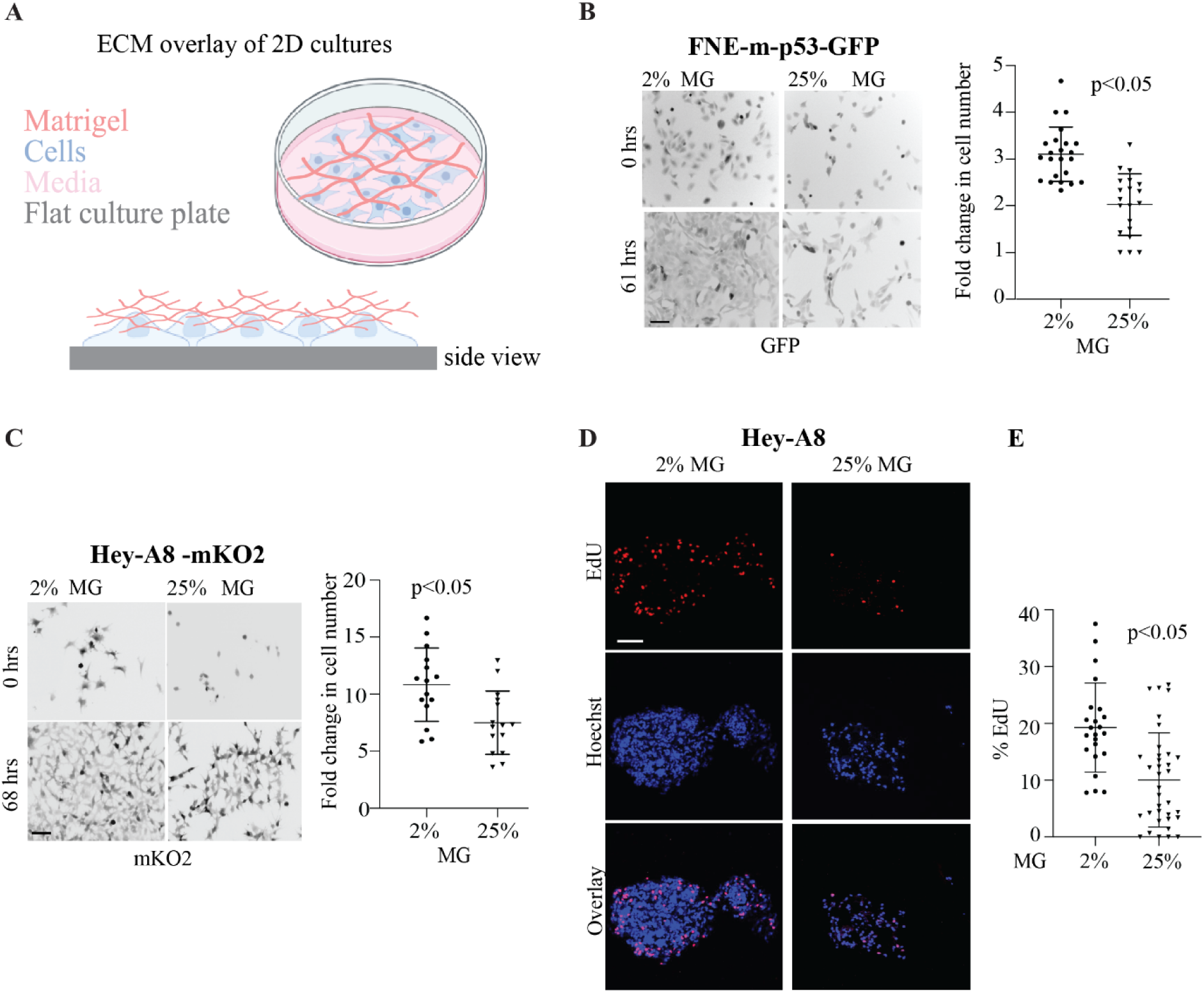
Elevation of ECM concentration suppresses FNE-m-p53 and Hey-A8 cell proliferation. (**A**) Cartoon representation of the experimental design for ECM overlay of cell monolayers. Created with BioRender.com. Representative fluorescent images of (**B**) FNE-m-p53 expressing GFP or (**C**) Hey-A8 cells expressing mKO2 monolayers overlayed with 2% or 25% MG. Dot plots represent a fold-change increase in cell number over time. Each dot corresponds to one field of view (FOV) containing (2-116 cells) and (3-108 cells) in B and C, with a total of 15-20 FOVs/ condition (2% & 25%). Data points are presented as mean ± SD. An unpaired, two-tailed, parametric student t-test with Welch’s correction was used to compute the statistical difference. (**D**) Representative fluorescent images of EdU and Hoechst in Hey-A8 spheroids reconstituted with 2% and 25% MG. Spheroids were processed 9 days after starting the culture. (**E**) The dot plot represents the fraction of EdU-positive cells. Each dot represents a spheroid; total spheroids 23 and 34 in 2% and 25%, respectively. All data points are shown with bars indicating mean ± SD. An unpaired, two-tailed, non-parametric, Mann-Whitney test was used to compute the statistical difference. Scale bars are 100 μm.

**S. Figure 4:**
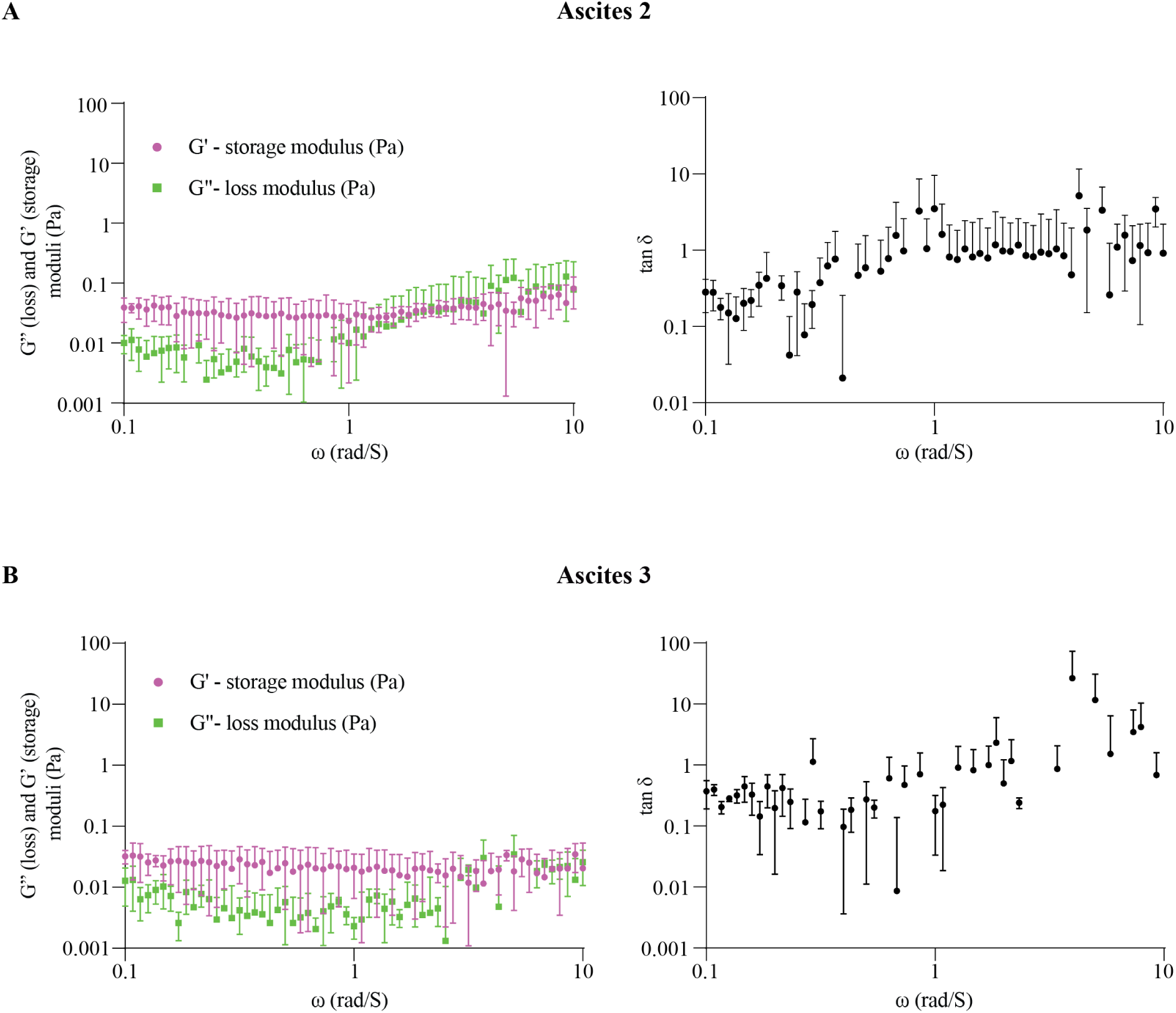
Viscoelastic properties of ascitic fluid samples isolated from OC patients with relapsed disease. Graphs represent the measurements of storage moduli (elastic), loss moduli (viscous), and tan δ (loss moduli divided by storage moduli) of (**A&B**) patient-derived ascites samples. Each data point represents the mean and SD from triplicate measurements with a total of 61 measurements.

**S. Figure 5.**
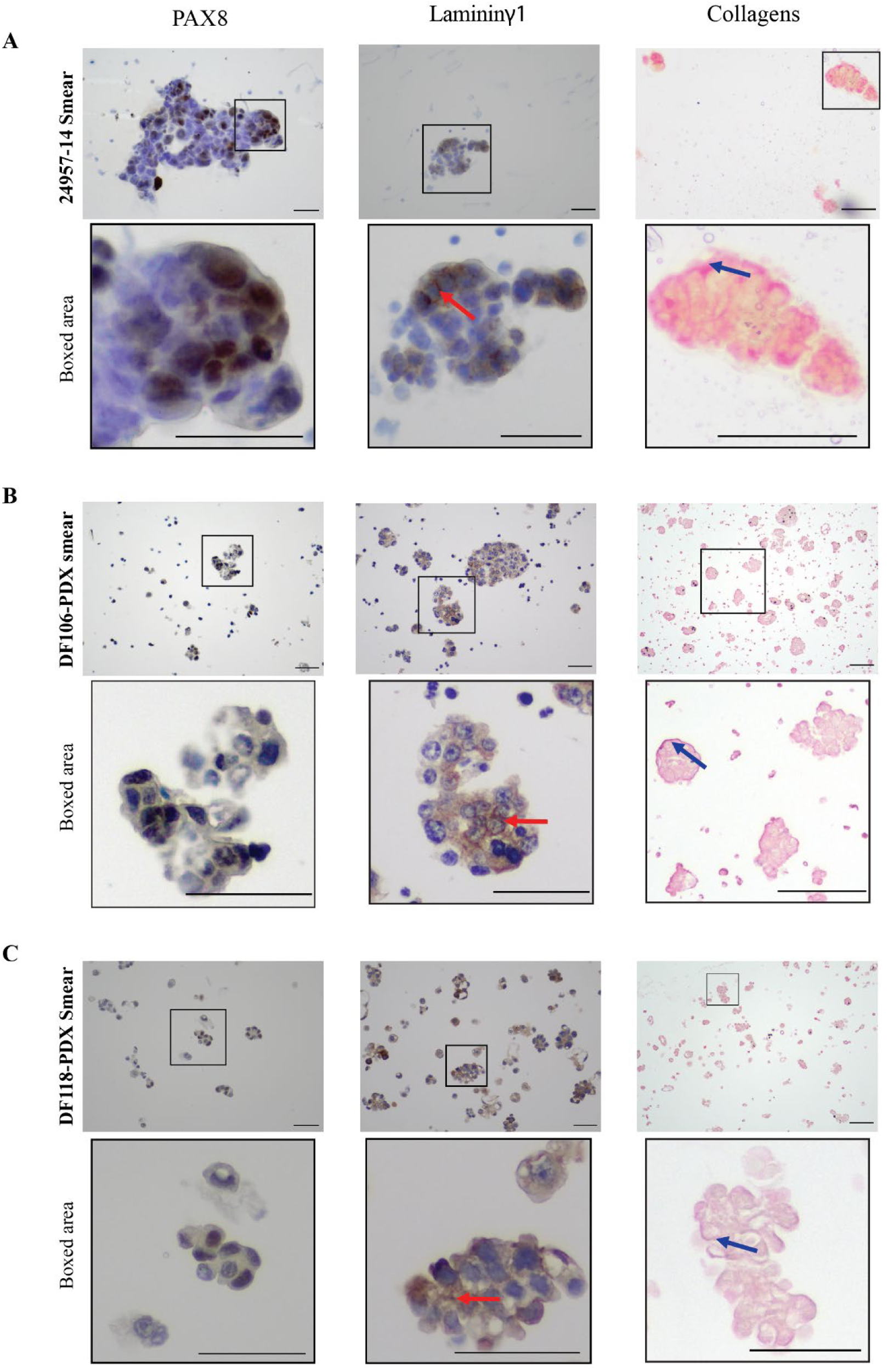
Laminin γ1 and collagen expression in OC clusters isolated from human or mouse ascites. Immunocytochemistry of Laminin γ1 and collagen in PAX8-positive (**A**) human, (**B & C**) PDX ascitic smears. Red arrows point to lamininγ1. Blue arrows point to collagen. Scale bars are 50 µm.

**S. Figure 6.**
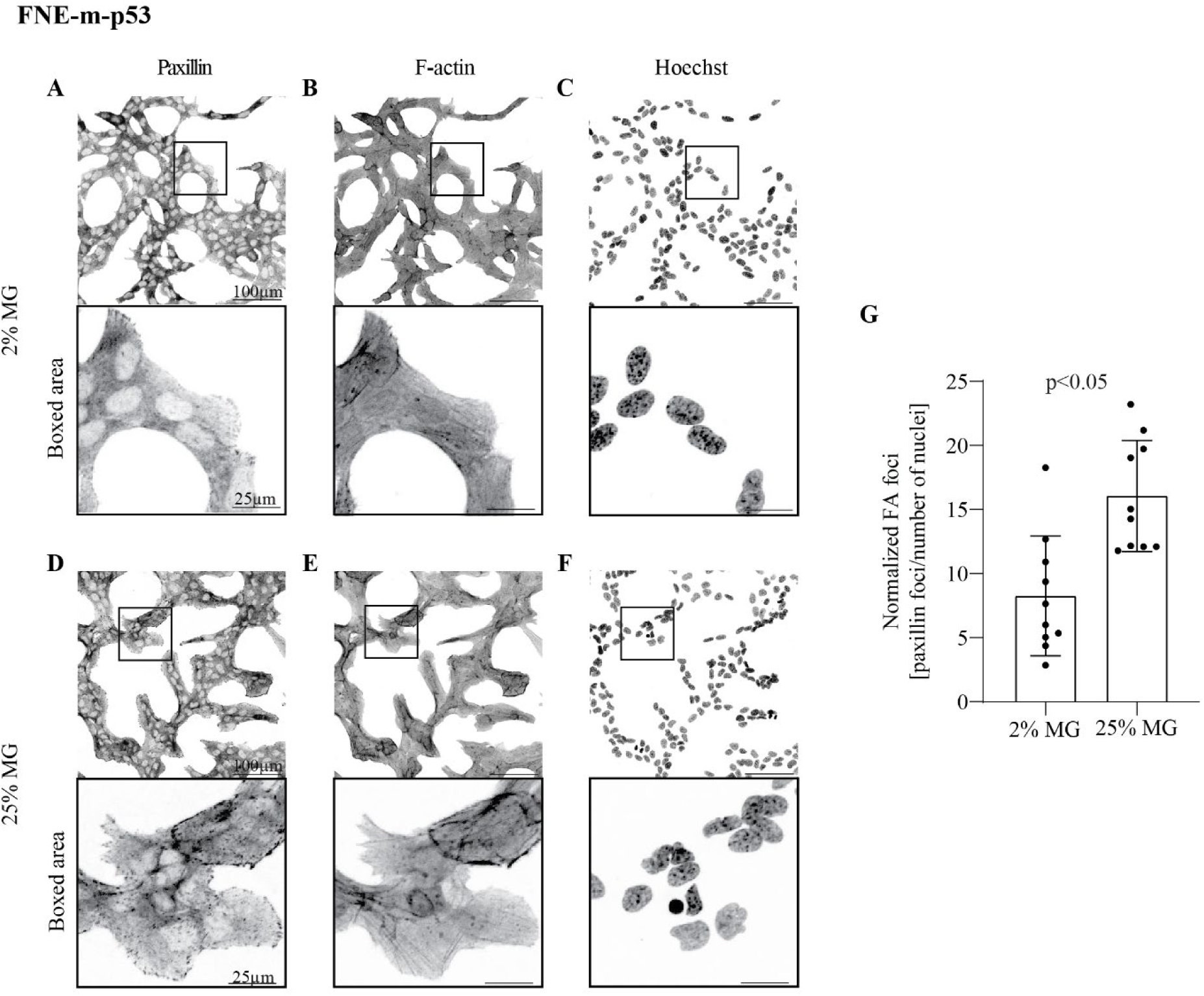
Evaluation of focal adhesions in FNE-mp53 cells overlayed with 2% or 25% MG. (**A**) Adhered monolayer of FNE-m-p53 cells was overlayed with (**A, B, C**) 2% or (**D, E, F**) 25% MG for 72 hours, fixed and stained by immunofluorescence with (**A, D**) antibodies against paxillin, (**B, E**) phalloidin-FITC (binding to F-actin), and (**C, F**) Hoechst 33342. (**G**) Quantification of the number of focal adhesions observed in FNE-m-p53 cells overlayed with 2% or 25% MG, respectively. Each data point corresponds to one field of view (FOV) containing (31-376 cells), with a total of 10 FOVs/ condition (2% & 25%). All data points are shown with bars indicating mean ± SD. An unpaired, two-tailed, parametric student t-test with Welch’s correction was used to examine statistical differences between data sets. Y-axis represents the number of paxillin foci normalized to the number of nuclei per ROI.

**S. Figure 7.**
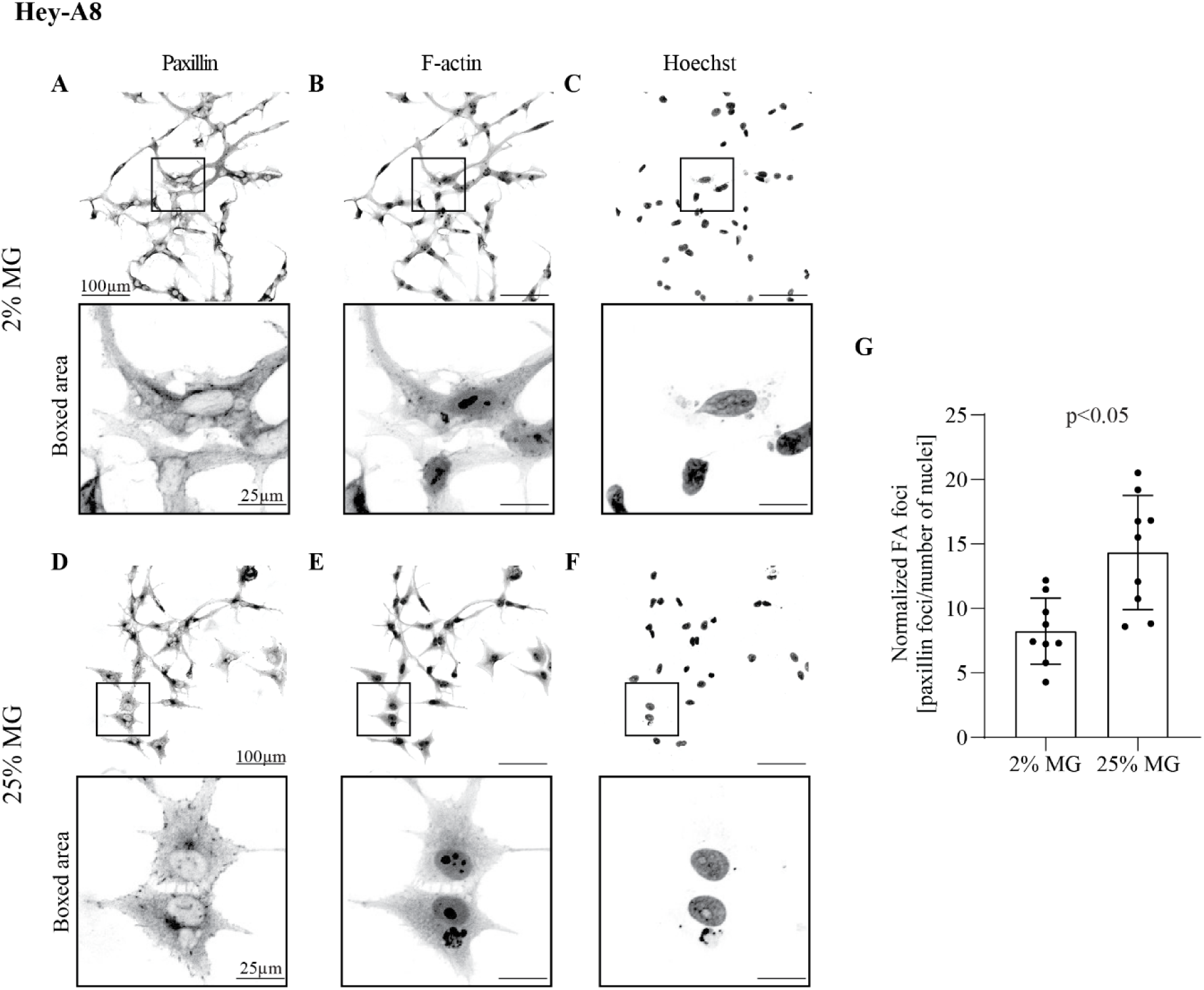
Evaluation of focal adhesions Hey-A8 monolayers overlayed with 2% or 25% MG. (**A**) Adhered monolayer of Hey-A8 cells was overlayed with (**A, B, C**) 2% or (**D, E, F**) 25% MG for 72 hours, fixed, and stained by immunofluorescence with (**A, D**) antibodies against paxillin, (**B, E**) phalloidin-FITC (binding to F-actin), and (**C, F**) Hoechst 33342. (**G**) Quantification of the number of focal adhesions observed in Hey-A8 cells overlayed with 2% or 25% MG, respectively. Each data point is one ROI (12-149 cells). All data points are shown with bars indicating mean ± SD. An unpaired, two-tailed, parametric student t-test with Welch’s correction was used to examine statistical differences between data sets. Y-axis represents the number of paxillin foci normalized to the number of nuclei per ROI.

**S. Figure 8.**
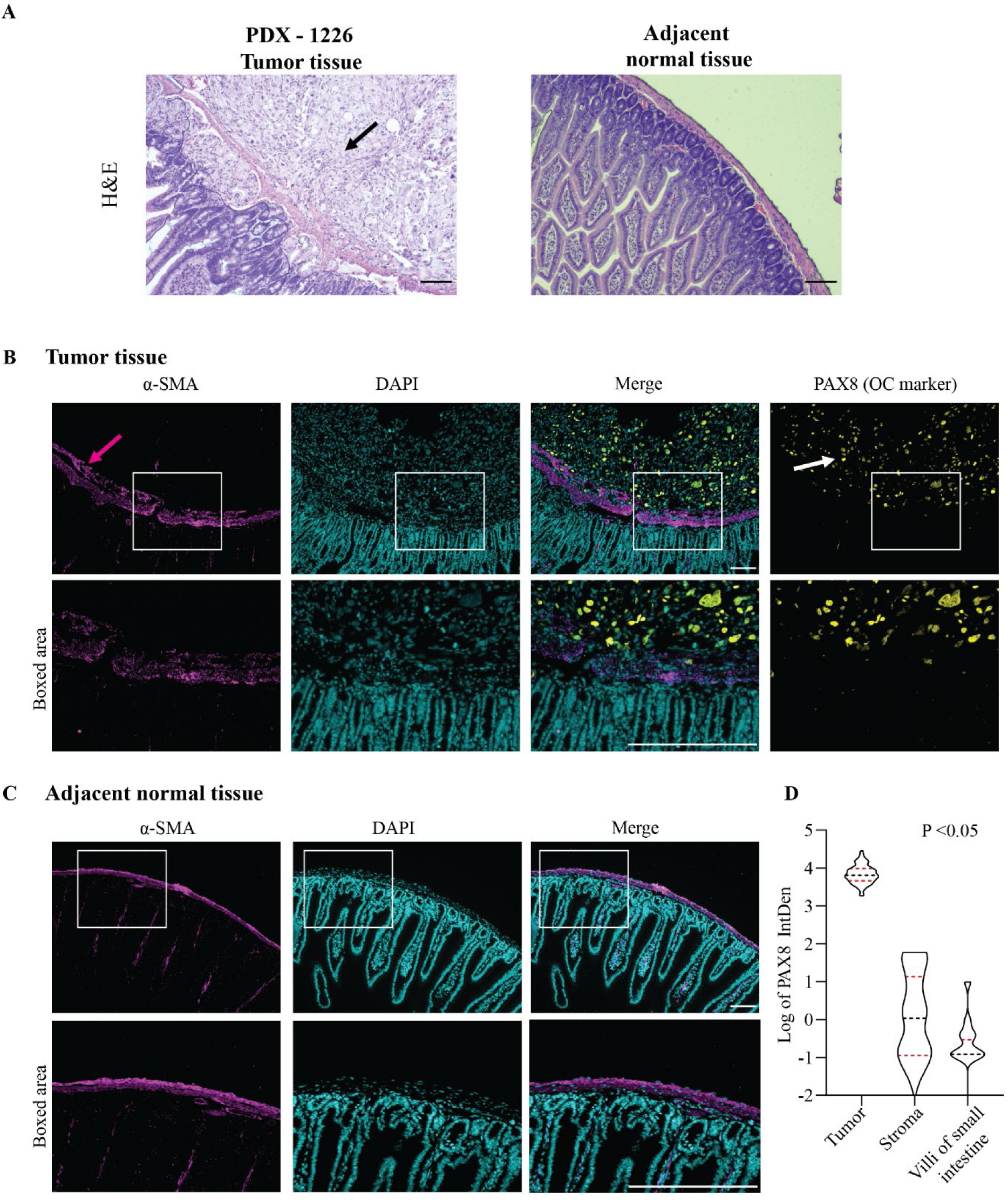
PDX-1226 growing tumor on the surface of the small intestine. (**A**) H&E of tumor nests (black arrow) and adjacent normal tissues. (**B**) Immunofluorescence of alpha smooth muscle actin (α-SMA) (magenta arrows), DAPI, PAX8 (white arrows) in tumor section. (**C**) Immunofluorescence of α-SMA and DAPI in adjacent normal tissue. Scale bars are 100 µm. (**D**) Quantification of PAX8 fluorescence intensity (integrated density). Each data point represents one region of interest (ROI). Total ROIs: tumor 66, stroma 20, and villi 20. All data points are shown as median and quartiles. An ordinary one-way ANOVA test was used to examine statistical differences between data sets. ANOVA was followed by Tukey post-hoc analysis; Tumor vs. Stroma P < 0.001, Tumor vs. Villi small intestine P < 0.001, Stroma vs. Villi small intestine P > 0.999.

## Supplementary Movies

**MOVIE 1.** Outgrowth formation by FNE-m-p53 spheroids reconstituted with 2% MG. Cell clusters were imaged for 5 days at 2-hr intervals.

**MOVIE 2.** Outgrowth formation by Hey-A8 spheroids reconstituted with 2% MG. Cell clusters were imaged for 5 days at 2-hr intervals.

**MOVIE 3.** Outgrowth formation by Tyk-nu spheroids reconstituted with 2% MG. Cell clusters were imaged for 5 days at 2-hr intervals.

**MOVIE 4**. FNE-m-p53-GFP cell proliferation in 2% and 25% MG. Images were obtained every 22 minutes for a duration of 61 hrs.

**MOVIE 5.** Hey-A8-mKO2 cell proliferation in 2% and 25% MG. Images were obtained every 22 minutes for a duration of 68 hrs.

**MOVIE 6.** Evolution of trajectories made by FNE-m-p53-GFP monolayer cultures overlayed with 2% or 25% MG. Images were obtained every 20 minutes for a duration of 61 hours. First frame of the movie corresponds to 13-hour time point.

**MOVIE 7.** Evolution of cell movement trajectories in Hey-A8-mKO2 monolayer cultures overlayed with 2% or 25% MG. Images were obtained every 15 minutes for a duration of 68 hours. First frame of the movie corresponds to 20-hour time point.

